# Combined heat and drought stress alters chromatin accessibility and primes potato abscisic acid biosynthesis

**DOI:** 10.64898/2026.11.11.765576

**Authors:** Tim Godec, Maja Križnik, Maja Zagorščak, Katja Stare, Živa Ramšak, Sophia Sonnewald, Lamis Abdelhakim, Klára Panzarová, Natalia Rodriguez Granados, Robert D Hancock, Mark A Taylor, Raymond Campbell, Rashmi Sasidharan, Salomé Prat, Maroof Ahmed Shaikh, Markus Teige, Marko Petek, Kristina Gruden

## Abstract

Due to climate change, abiotic stresses get increasingly more frequent, in a combined and recurrent manner in the same growing season. Chromatin accessibility enables efficient responses to combined stress and priming for reoccurring stresses. To study the regulatory basis of priming, we profiled chromatin accessibility and gene expression in leaves of two genetically distinct potato cultivars after prolonged heat, drought, waterlogging, and combined heat and drought. We show that most accessibility changes are stress-specific, however, drought stress elicits a distinct transcriptional response as compared to combined heat and drought stress, with strong chaperone induction and dampened abscisic acid signalling, despite water deficit. A second cycle of combined stress reveals transcriptional memory of abscisic acid biosynthesis gene *StNCED3* and its transcriptional activator StWRKY33b, providing a new mechanistic insight of abiotic stress memory in a staple crop important for global food security.

## INTRODUCTION

Potato (*Solanum tuberosum*) is one of the world’s most important food crops and a key contributor to global food security (Devaux et al., 2020). Yet, potato is particularly vulnerable to abiotic stress, including heat, drought and waterlogging, which can disrupt plant growth and tuber development and reduce yield (Zagorščak et al., 2025). Climate change is expected to increase the frequency and severity of such stresses, making improved crop resilience an important breeding target (Zandalinas & Mittler, 2022). These stresses rarely occur in isolation under field conditions; periods of high temperature often coincide with declining soil water availability, and drought or heat may follow periods of excessive rainfall (Zhang & Sonnewald, 2017). This creates a need to understand plant responses not only to individual stresses, but also to their occurrence in combination and sequence (Zagorščak et al., 2025). Responses to combined or successive stress cannot necessarily be inferred from responses to each individual component (Jiang et al., 2025). In *Arabidopsis*, combined heat and drought produce distinct physiological and transcriptional responses specific to the stress combination (Jiang et al., 2026; Morales et al., 2022; Rizhsky et al., 2004). Similar combination-specific responses have been observed across multifactorial and sequential stress treatments in other plants (Jiang et al., 2026; Zandalinas et al., 2021; Zandalinas & Mittler, 2022; Zhang & Sonnewald, 2017), suggesting that interactions among stress signals can generate regulatory states that are not captured by single-stress conditions. However, the regulatory basis of these combination-specific transcriptional responses remains poorly understood.

Chromatin accessibility represents an important regulatory layer underlying transcriptional responses to environmental stress. Regulatory elements such as promoters and enhancers must be accessible to transcription factors and the transcriptional machinery, and changes in chromatin accessibility can therefore alter the potential for gene regulation (Lu et al., 2019). Assay for transposase-accessible chromatin using sequencing (ATAC-seq) enables genome-wide profiling of these accessible regions (Buenrostro et al., 2013). ATAC-seq offers a broader regulatory read-out than other epigenetic approaches because it directly identifies regions of accessible chromatin, whereas DNA methylation and histone-based assays measure additional regulatory features. A study in Arabidopsis mapping accessible chromatin identified thousands of environmentally responsive regulatory elements and enabled reconstruction of the transcription factor circuits underlying the heat-shock response (Sullivan et al., 2014). Similar coupling between accessibility and expression has been reported under heat and waterlogging in rice (Liang et al., 2021; Reynoso et al., 2019), and across heat, salt and drought in the xerophyte *Zygophyllum xanthoxylum* (T.-Q. Wang et al., 2025). In tomato, integrated ATAC-seq and RNA-seq analysis identified differential accessibility associated with natural variation in the promoter of *SlWRKY34* affecting cold tolerance (Guo et al., 2024). These studies illustrate the link between regulatory variation and chromatin accessibility in stress, while regulation of plant acclimatisation to combined stress and priming for reoccurring stress conditions is less well understood (Gallusci et al., 2023). Priming is defined as the enhanced capacity to respond to a reoccurring stress following an initial stress exposure, mediated by stress memory (H. Liu et al., 2022). One mechanism explaining this long-term (memory) phenomenon is that stress is imprinted in changes of chromatin organization. For example, it was shown that DNA methylation patterns play an important role in immune priming in Arabidopsis (Kurtz et al., 2025; Parker et al., 2025). Transcriptional signatures of abiotic stress priming in potato remain poorly characterised, although mild heat priming has been shown to alter transcript dynamics following subsequent heat shock, with chromatin remodelling among the implied mechanisms (Trapero-Mozos et al., 2018).

Responses at the chromatin level after prolonged abiotic stresses and their combinations in potato remain largely unexplored. Existing potato stress-treatment chromatin accessibility studies have focused on cold acclimation in a potato wild relative, or drought and salt treatments (X. Li et al., 2024; Wen et al., 2025) in the potato DM genotype (Wen et al., 2025). Recent analysis of two contrasting cultivars found substantial genotype-dependent DNA methylation differences, but limited heat-induced methylation changes, and no consistent correspondence between differentially methylated regions and heat-responsive gene expression (Yeo et al., 2026). Moreover, to our knowledge, chromatin accessibility under waterlogging stress, or under combined or sequential heat and drought, has not yet been investigated at the genome-wide level in potato.

Here, we address this gap by profiling chromatin accessibility with ATAC-seq in leaves of two genetically distinct potato cultivars, Désirée and Lady Rosetta, subjected to prolonged heat, drought and waterlogging, as well as to a combined heat and drought stress. RNA-seq was used to complement the accessibility data and identify transcriptional responses associated with chromatin remodelling. Because heat and drought frequently recur and coincide within a single growing season (Zhang & Sonnewald, 2017), this combination is the most relevant for studying plant responses to repeated stress. To test whether the observed responses reflect a priming effect, we exposed Désirée, the cultivar with the stronger chromatin response, to two cycles of combined drought and heat and followed its transcriptional response. Using this approach, we identified novel regulators of abscisic acid (ABA) biosynthesis in plants carrying a memory of the first stress.

## RESULTS

### Prolonged abiotic stresses induce distinctive changes in the accessible chromatin landscape of potato leaves

Plants that have experienced stress are better prepared to withstand the reoccurring adverse conditions. Here we asked whether diverse abiotic stresses leave an imprint in chromatin organisation, which can determine potato acclimation to upcoming abiotic stress. We exposed mature plants to prolonged heat, drought or waterlogging, and after heat followed by combined heat and drought (heat–drought), designed to reflect the stress progression under climate change conditions (Fig. 1A). To capture universal chromatin structure rearrangement across different potato genotypes, we analysed two cultivars, Désirée and Lady Rosetta, which have distinct breeding histories and therefore distinct genomic backgrounds (van Berloo et al., 2007). Across the experiment we identified ∼147,000 accessible chromatin regions (ACR) covering ∼6% of the potato genome. ACRs were detected in promoters of over 80% of transcribed genes, indicating high sensitivity of the applied methodology (Suppl. Fig. S1). ACRs were enriched around the transcription start sites (TSS) (Suppl. Fig. S2B). Nearly one third of ACRs were distal intergenic (located more than 3 kb from TSS), confirming that regulatory regions extend well beyond proximal promoters (Suppl. Fig. S2C).

**Figure 1.**
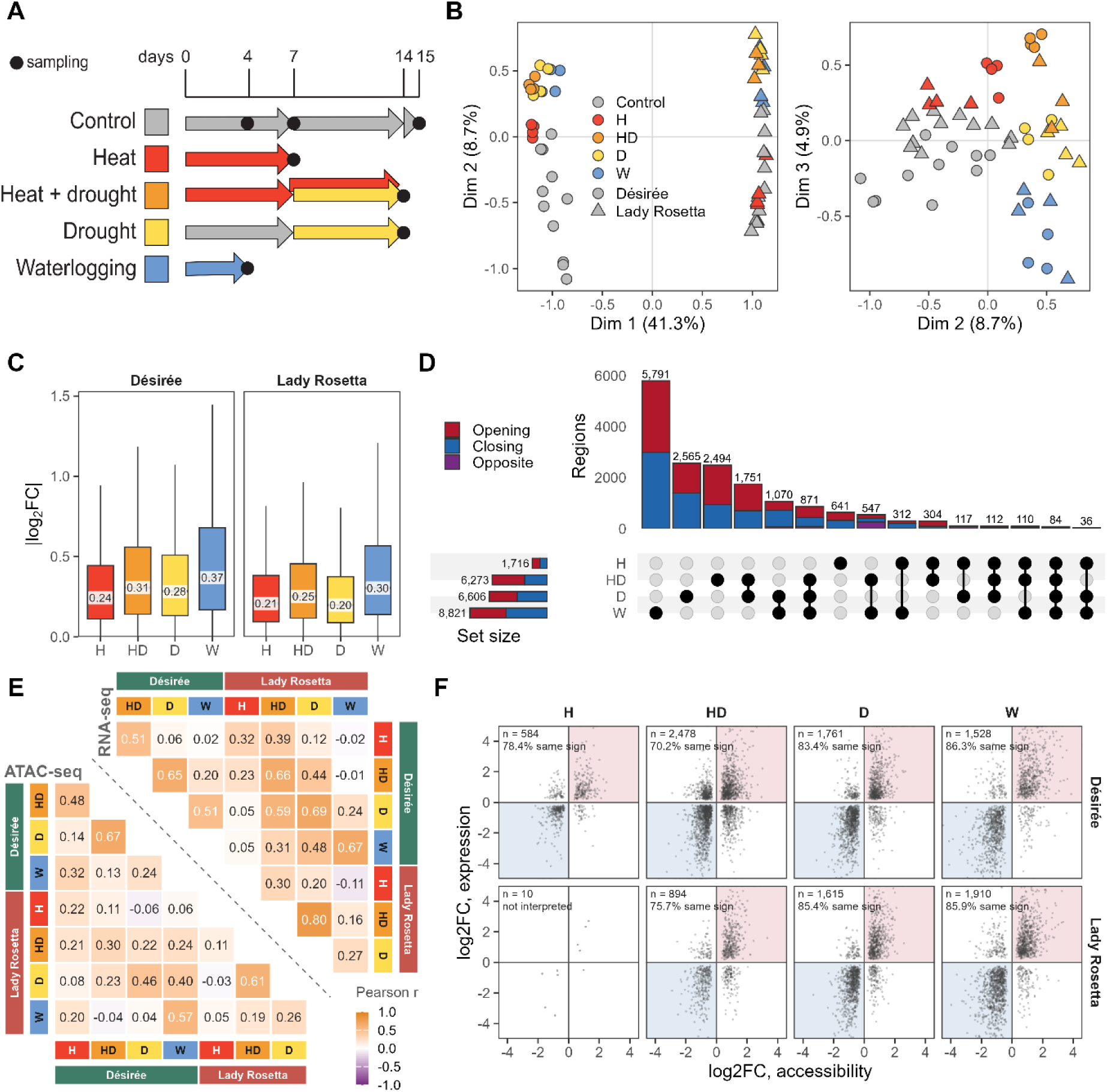
Accessible chromatin landscape of potato leaves in response to diverse abiotic stresses. **A)** Experimental design of initial abiotic stress experiment showing stress treatments, treatment durations, and sampling time points. **B)** Sample diversity based on detected accessible chromatin regions: Multidimensional scaling (MDS) of the filtered peaks across samples. MDS dimensions 1 and 2 (left) and dimensions 2 and 3 (right) are shown. Percentages of variation are given in parentheses. **C)** Distribution of absolute log₂ fold changes (|log₂FC|) across 56,724 filtered peaks for each stress and cultivar. Boxes indicate the median and interquartile range (IQR), and whiskers extend to 1.5 × IQR. **D)** Overlap of differentially accessible regions (|log₂FC| > 0.5, FDR < 0.05; response averaged over both cultivars) among treatments. **E)** Pearson correlations of log₂ fold changes for accessible regions (lower left) and expressed genes (upper right) between different stress treatments. **F)** Relationship between promoter accessibility and transcript log₂ fold change for genes showing significant responses in both assays (FDR < 0.05), shown separately by cultivar and stress. For each gene, the promoter region with the lowest FDR in the differential accessibility analysis was selected as the representative region. Shaded quadrants indicate concordant responses, and the percentage of genes showing concordant responses (i.e., changes with the same sign in both assays) is indicated in each plot. n denotes the number of genes. ATAC-seq, assay for transposase-accessible chromatin with sequencing; FDR, false discovery rate; H, heat; D, drought; W, waterlogging; HD, combined heat and drought.

We compared the four regimes to test whether chromatin reorganization is shared or specific for the stress responses in our study. Waterlogging caused the largest accessibility changes in both cultivars, followed by heat–drought and drought. (Fig. 1C). Heat had the weakest effect in Désirée, whereas Lady Rosetta heat-treated plants were barely distinguishable from controls. Beyond differing in magnitude, the accessibility changes were also highly stress-specific: about two thirds of responsive regions responded to only one regime, and fewer than a hundred responded to all four (Fig. 1D). The largest shared sets involved drought, together with heat–drought, waterlogging, or both. The overlap with heat–drought is expected from their shared water deficit, whereas the overlap with waterlogging suggests it shares part of the drought response. Regions that opened under drought, heat–drought and waterlogging lay at canonical ABA- and dehydration-responsive genes, among them genes encoding ABF2, a HAI-type clade A protein phosphatase 2C (PP2C), the dehydrin RD29B, an HD-Zip I homologue of AtHB7 and AtHB12, and the trehalose-6-phosphate synthase TPS11 (Suppl. Table S5). The accessibility response is therefore largely stress-specific, but drought, heat– drought, and waterlogging converge on a shared set of ABA-responsive regulatory regions.

Given their different breeding histories, we compared the chromatin landscapes of the two cultivars and their responses to stress. Both had a similar number of accessible regions (127,196 in Désirée and 124,383 in Lady Rosetta; Suppl. Table S1), but the accessibility of these regions differed, separating cultivars more strongly than stress treatments (Fig. 1B). In response to stress, the cultivar differences were most pronounced in heat–drought and much smaller in drought and heat, while waterlogging reshaped almost the same regions in both (Suppl. Table S2).

We next compared those chromatin changes with the transcriptional response profiled in the same experiment. Among genes that responded on both molecular layers, promoters tended to open at induced genes and close at repressed genes consistently across cultivars and stress treatments (Fig. 1F). However, the magnitude of accessibility changes correlated onlyb moderately with the expression changes observed across the genes with a detected promoter peak (Suppl. Fig. S4A), and differentially accessible promoters was accompanied with fewer than one in five of the differentially expressed genes with a promoter peak (Suppl. Fig. S4B). Accessibility changes therefore indicate the direction, but not the extent, of the transcriptional response.

### Abiotic stress reshapes chromatin accessibility across photosynthesis, growth and defence related processes

Having established that accessibility changes track the direction of transcriptional responses, we next asked which biological processes these changes affect. Stress reduced accessibility of photosynthetic, plastid and growth-related genes, and increased accessibility of defence-associated pathways (Fig. 2A, Suppl. Fig. S5, Suppl. Table S3). Under heat and waterlogging, promoter accessibility of photosynthesis genes decreased strongly and coherently in both cultivars, across most photosynthetic subcategories, and gene expression followed the same trend (Fig. 2A, Suppl. Table S3). Under drought, the cultivars responded in opposite directions: in Désirée, both accessibility and expression of photosynthesis genes increased strongly, whereas in Lady Rosetta both decreased, with significant accessibility changes seen only for photosynthesis as a whole, photophosphorylation and for chlororespiration. Under heat– drought, accessibility of photosynthesis-related genes did not change significantly in either cultivar, apart from a decrease in photophosphorylation and photosystem II-related genes in Lady Rosetta, although their expression decreased, less than under heat or waterlogging. Under waterlogging, the response in both cultivars extended beyond photosynthesis to broader plastid-related processes, including chlorophyll metabolism, plastid gene expression and organellar translation, together with reduced accessibility of chloroplast protein import pathways and, in Désirée under waterlogging, of mitochondrial complex I (Suppl. Table S3). Additionally, the enrichments indicate reduced accessibility across several components of the cell-growth machinery, spanning signalling, cytoskeletal organisation, cell-wall organisation and cell division (Fig. 2A, Suppl. Fig. S5). The cell-division response involved both cyclins and cell-cycle inhibitors, indicating reduced accessibility of genes involved in the cell-cycle biological process, as a whole.

**Figure 2.**
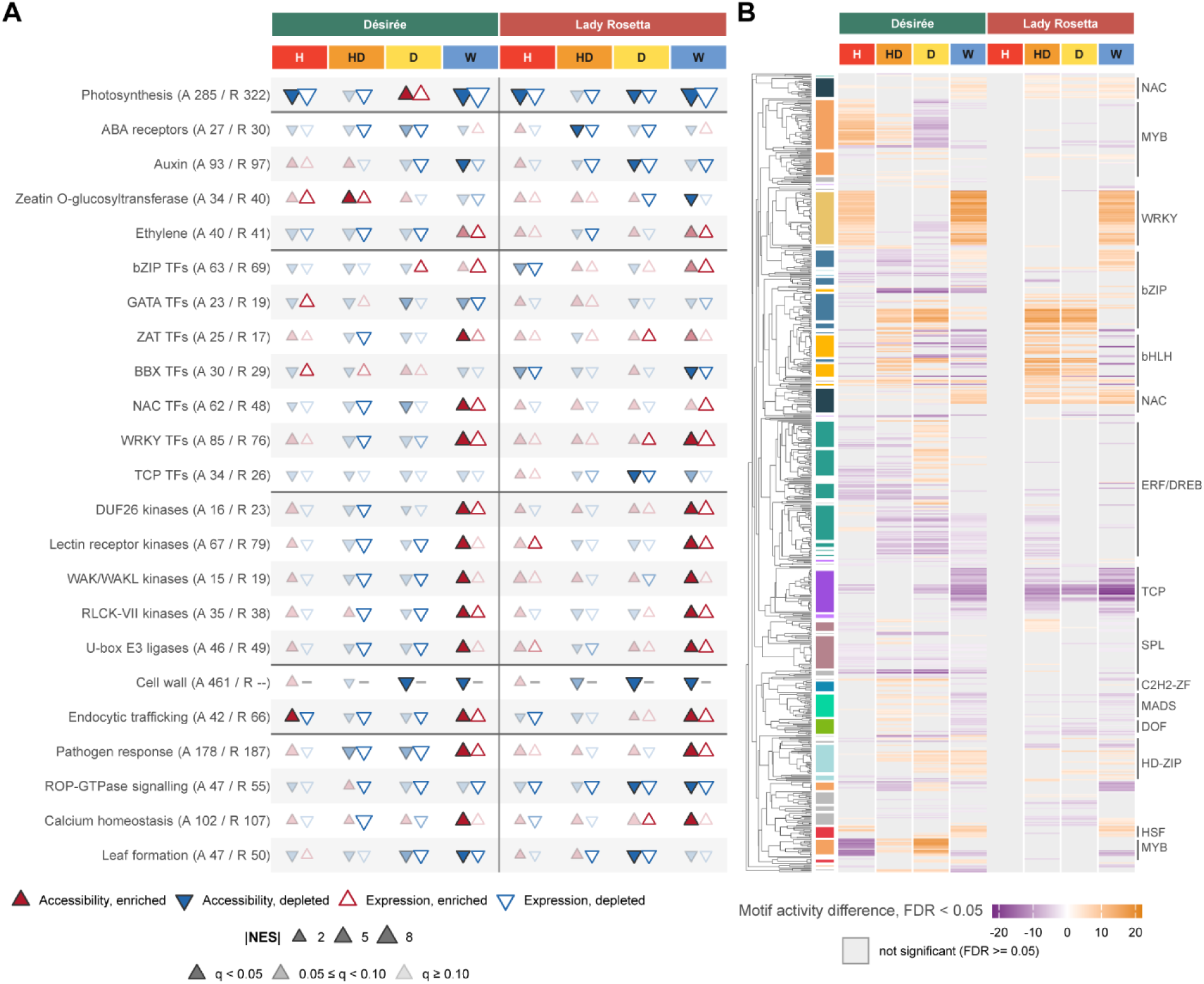
Process level integration analysis of chromatin accessibility and gene expression. **A)** Enrichment of MapMan gene sets across all comparisons. Filled and open triangles indicate enrichment in the accessibility and expression datasets, respectively; triangle orientation indicates increased or decreased accessibility or expression, and triangle size represents absolute value of the normalized enrichment score (NES). Opacity of triangles indicates q value: <0.05, 0.05–0.10, and >0.10 correspond to 1, 0.5, and 0.22, respectively. Selected MapMan gene sets are shown, grouped by process and separated by horizontal lines; all 45 gene sets are shown in Suppl. Fig. S5. Full results of enrichment analysis are available in Suppl. Table S3. In parenthesis: number of genes in the gene set for accessibility (A) and expression (R) datasets. **B)** Differential activity of transcription factor motifs (i.e. relative change in chromatin accessibility of ACRs containing the motif) across all comparisons. Motifs were obtained from the JASPAR 2026 plants CORE collection. Cell colour indicates differences in motif activity at FDR < 0.05 (orange – higher activity, violet – lower activity); grey cells are not significant. Rows are ordered according to an average-linkage hierarchical clustering based on 1 − r, where r is the Pearson correlation coefficient derived from the RSAT pairwise motif-similarity matrix. Coloured boxes (left column) indicate transcription factor families; grey was used for small families.

In contrast, defence-associated pathways gained promoter accessibility, most prominently in waterlogging (Fig. 2A). In both cultivars, waterlogging increased accessibility of the pathogen, effector-triggered immunity (ETI), TNL-mediated ETI and defence-mechanism pathways, accompanied by increased transcript abundance. This agrees with flooding- and hypoxia-induced changes in the transcriptomes, in which WRKY transcription factors and innate-immunity genes are induced (Hsu et al., 2013; Tamura & Bono, 2022). Waterlogging also increased accessibility of G-type lectin receptor-like kinases, which were concentrated in tandem genomic clusters.

Transcription-factor families also changed in accessibility, mostly in a stress-specific way (Fig. 2A). Under waterlogging, WRKY genes gained accessibility in both cultivars, NAC genes in Désirée, and ZAT genes in Désirée and, although weaker, also in Lady Rosetta (q < 0.10). TCP genes showed a depletion that was consistent in direction across most comparisons but significant only in Lady Rosetta under drought, and under waterlogging. BBX genes lost accessibility in Lady Rosetta under waterlogging, and GATA genes, to a lesser extent, in Désirée under drought and waterlogging (q < 0.10). bZIP genes changed little (q < 0.10 only in Lady Rosetta, under heat and waterlogging; Fig. 2A). Inferred motif activity in ACRs supported the changes in WRKY, NAC, TCP and GATA, whose motifs changed in the same direction in the corresponding stresses (Fig. 2B), but not in ZAT, whose C2H2 zinc-finger motifs did not gain activity under waterlogging. BBX could not be assessed because this family is not represented in the motif library.

Motif activity analysis additionally identified a strong gain of G-box/ABRE regulatory activity under drought and heat–drought, which was not captured among the leading GSEA categories (Fig. 2B, Suppl. Table S4). The ABRE-type bZIP archetype showed the strongest gain in motif activity under water deficit. (Suppl. Fig. S6). This motif was highly enriched in both drought and heat–drought, particularly in Lady Rosetta, but showed little or no response to heat or waterlogging. The ABRE response therefore represents a drought-specific chromatin signature rather than a general response to abiotic stress. We also checked for known heat response transcriptional regulation. Indeed, HSF motif enrichment was observed following heat in Désirée, but not in Lady Rosetta (Fig. 2B, Suppl. Table S4).

### Combined heat and drought stress induces a unique regulatory response

Combined stresses can elicit responses that are not predictable from the single stresses, but whether this extends to chromatin organization is unknown. We therefore asked how the combined heat–drought response relates to those of heat and drought alone. At the genome-wide level, heat–drought accessibility changes correlated more strongly with drought than with heat in both cultivars (Fig. 1E, Fig. 3A), and the same held true for expression changes (Fig. 3A). Rather than a modified drought response, heat–drought elicited a largely unique response. In the transcriptome, 2,327 genes in Désirée and 1,618 in Lady Rosetta responded specifically to heat–drought, without significant responses to either individual stress, forming the largest treatment-specific gene set in both cultivars (Fig. 3B). Comparison with the individual-stress responses further categorized genes according to how their responses were altered under heat–drought (Fig. 3A). Emergent responses were the largest class in both cultivars. In addition, many drought-responsive genes kept their direction under heat–drought but were amplified or attenuated.

**Figure 3.**
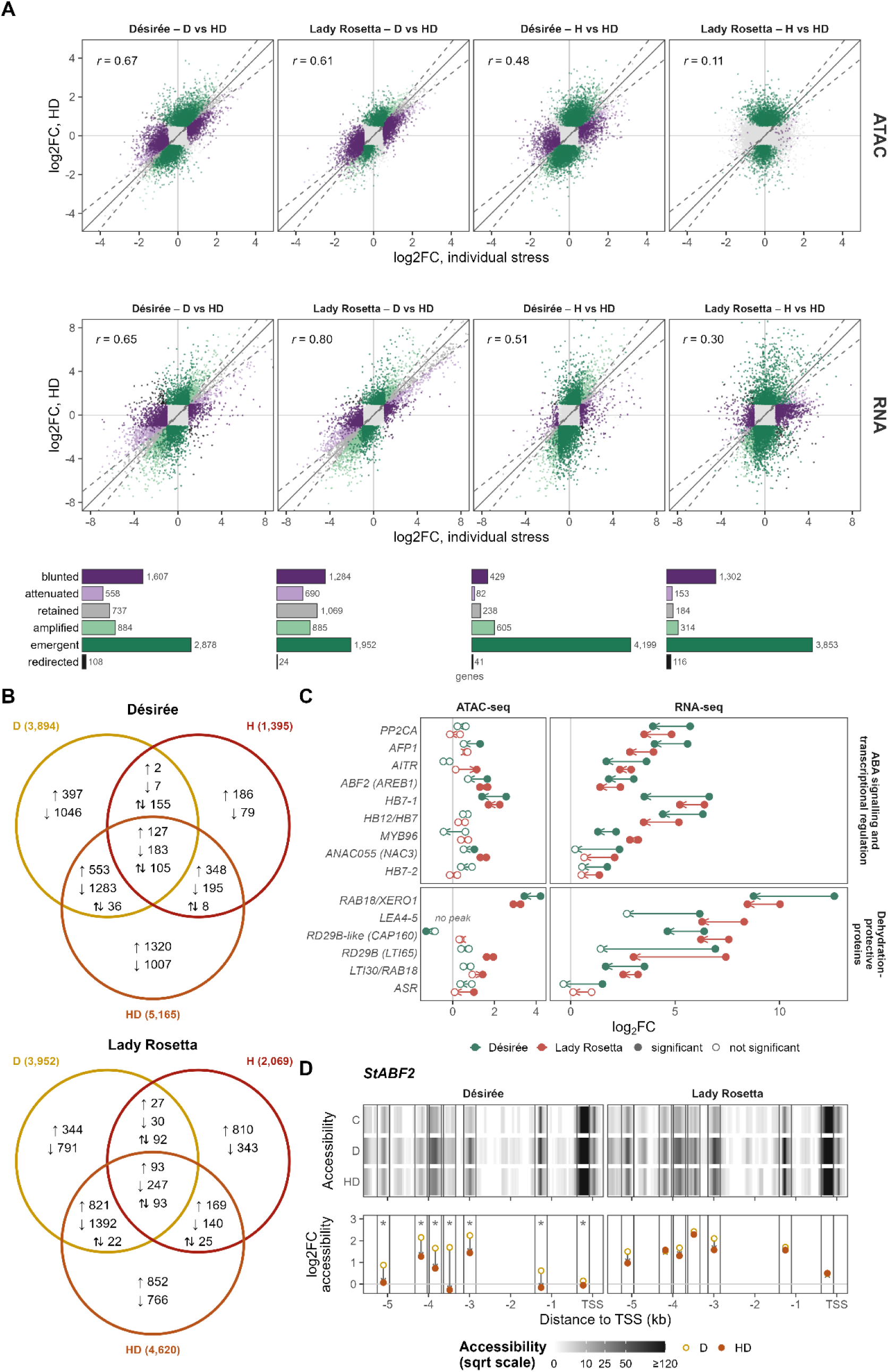
Combined heat and drought stress induces a largely unique transcriptional and chromatin accessibility response. **A)** Chromatin accessibility (top, one point per peak, n = 56,724) and transcript (bottom, one point per gene, n = 22,134) responses to combined heat and drought (HD), plotted against the individual heat (H) or drought (D) response in Désirée and Lady Rosetta. Grey points respond to neither stress (responsive = FDR < 0.05 and |log₂FC| > 0.5 for peaks, |log₂FC| > 1 for transcripts). Coloured points are classed by how they respond under heat–drought compared with the individual stress: *emergent* (responsive under heat–drought only), *blunted* (responsive to the individual stress only), *redirected* (opposite direction) or, when the direction is the same, *amplified*, *retained* or *attenuated* (the heat–drought log₂FC is more than 1.2 times, 0.8 to 1.2 times, or less than 0.8 times the individual-stress log₂FC; dashed lines). The bars below the scatter plots give the number of genes in each class and are the colour key for both rows. Pearson r over all peaks or genes is shown in each plot. **B)** Venn diagram showing overlap of differentially expressed genes (FDR < 0.05 and |log_2_FC| > 1) in individual heat and drought, and combined heat–drought stress in cv. Désirée (top) and cv. Lady Rosetta (bottom). Arrows show upregulated (up), downregulated, and disagreeing (up, down) gene overlaps. **C)** Expression and chromatin accessibility responses of 15 drought-induced genes, grouped as ABA signalling and transcriptional regulation and dehydration-protective proteins. Gene short names are those of the Arabidopsis orthologues. Beginning of the arrow indicates responses in drought, end of the arrow indicates responses in heat–drought; filled symbols, FDR < 0.05; open symbols, not significant; ’no peak’, no tested accessible region. **D)** The *StABF2* upstream region. Top: mean accessibility in control, drought and heat–drought (ATAC-seq coverage normalized to genome-wide depth, where 1 is the genome average; square-root colour scale, capped at 120). Bottom: accessibility log₂FC relative to control under drought (open) and heat–drought (filled) at each tested region; asterisks, FDR < 0.05 for the fitted heat–drought versus drought contrast. ATAC-seq, assay for transposase-accessible chromatin with sequencing; FDR, false discovery rate; TSS, transcription start site.

Functional enrichment supported this interpretation (Suppl. Fig. S7). Protein quality control was the most significant and most consistently intensified response under heat–drought in both cultivars, with the chaperone response approximately doubling in heat–drought compared to drought or heat (Suppl. Fig. S7). This increase was associated with stronger induction of representative HSP90, HSP101/HSA32/CLPB3 and HSP70/HSP40 chaperone components, consistent with an increased protein-folding burden under combined stress. Interestingly, the changes in chromatin accessibility were mild for *HSP* genes, albeit the HSF motifs activity was increased. Only individual *StHSP90* and *StHSP70* genes had increased chromatin accessibility in heat and drought, respectively (Suppl. Table S5).

In contrast, ABA signalling and dehydration-protective genes were more prominent among drought-induced genes whose response was attenuated or blunted under heat–drought, including potato orthologues of Arabidopsis *ABF2*, *AFP1*, *PP2CA*, *MYB96* and *RD29B* (Fig. 3C). Changes in their chromatin accessibility were smaller and less consistent under combined stress. At the *StABF2* locus, drought-induced accessibility at distal upstream regions was selectively reduced under heat–drought. Accessibility 1–5 kb upstream of the TSS decreased in the same direction in both cultivars, significantly at all six upstream regions in Désirée, whereas the TSS-proximal region changed only slightly (Fig. 3D). Thus, the reduced StABF2 expression response under heat–drought is associated with attenuation of accessibility mainly at upstream regions rather than at the TSS-proximal region. Independent evidence from cold-acclimated *Solanum commersonii* further supports StABF2 as an abiotic-stress regulator (X. Li et al., 2024). We therefore prioritised StABF2 as the leading candidate associated with the water-deficit-induced gain in ABRE accessibility.

### Abscisic acid synthesis is in a heat–drought stress primed state brought by StWRKY33b

We next asked whether chromatin accessibility contributes to stress memory, helping the plant to respond more effectively to a recurring cycle of adverse conditions. We performed an additional independent experiment on the more responsive cv. Désirée, exposing plants to heat, drought and heat–drought for 4 days, then the drought-exposed plants were re-watered and exposed to the second round of drought for 4 days (Fig. 4A). As heat alone did not leave a substantial mark in chromatin accessibility (Fig. 1C, D, Suppl. Fig. S3), we focused on elucidating drought- and heat–drought-inflicted priming (sample description in Suppl. Table S6; soil moisture and plant physiology in Suppl. methods and results, Suppl. Tables S7, S8 and Suppl. Fig. S8, S9). We performed transcriptomics analysis at the beginning and end of each stress cycle.

**Figure 4.**
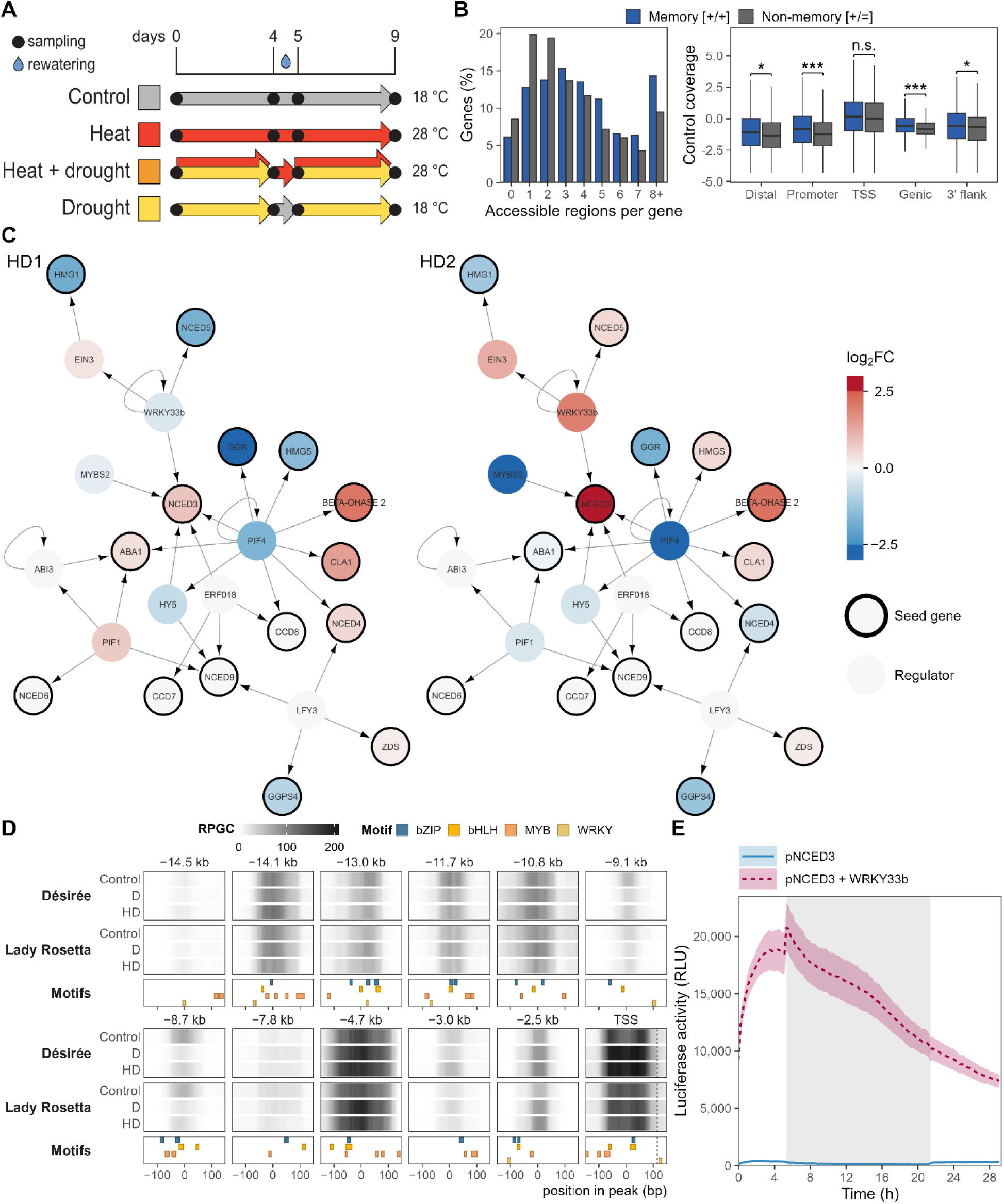
Priming of plants for additional cycles of drought and heat stress through ABA synthesis is modulated by StWRKY33b. **A)** Experimental design of priming experiment: 18 °C, 28 °C: temperature at which plants were grown; blue drop: time point of rewatering of drought- and heat and drought-stressed plants; black dots: tissue sampling points. **B)** Chromatin accessibility of drought-memory genes in control Désirée plants. Memory genes (induced in the first drought and more strongly in the second) are compared with non-memory genes induced equally in both droughts. Left: number of accessible regions per gene (test in Suppl. Fig. S10). Right: mean accessibility in control plants at five positions: distal upstream (3–10 kb upstream of the TSS), promoter (0.5–3 kb upstream), TSS-proximal (TSS ± 500 bp), gene body and 3′ flank (1 kb downstream of the gene end). Accessibility is expressed as the ATAC-seq coverage scaled to the mean coverage of all TSS regions (log₂ scale). Asterisks, Wilcoxon rank-sum test, Holm-adjusted (*, P < 0.05; **, P < 0.01; ***, P < 0.001; n.s., not significant). **C)** Network analysis of ABA synthesis genes: First neighbours of isoprenoid/ABA synthesis genes (seed list, dark black border) were extracted from a Comprehensive knowledge network (Bleker et al., 2024). The extracted network was further reduced to retain only transcriptional regulation edges to identify the transcription factors involved in regulation of ABA synthesis and gene expression data were mapped onto the network. Node fill colour denotes expression log₂ fold changes (log₂FC) in drought and heat-stressed vs control plants (HD1, left panel) and in re-droughted (after recovery) heat-stressed vs control plants (HD2, right panel). Red – higher expression in stress compared to control, blue – lower expression in stress compared to control. Arrows indicate direction of interaction. For full names and IDs of all genes in the network see Suppl. Table S12. **D)** Accessibility of *StNCED3* upstream region. Columns show accessible regions. Top two boxes: mean chromatin accessibility across treatments measured as ATAC-seq coverage normalized to genome-wide depth as reads per genomic content (RPGC). Bottom box: positions of bZIP (blue), bHLH (amber), MYB (orange) and WRKY (pale yellow) transcription factor binding motifs. **E)** Transcriptional regulation of *StNCED3* by StWRKY33b was confirmed *in planta* by transactivation assay in *Nicotiana benthamiana*. The luciferase activity of pStNCED3::iLUC was observed through relative luminescence units (RLU) measured at 30-minute intervals. The pStNCED3::iLUC construct alone served as control of basal activity. Median values with standard error of median for 16 replicates are reported. Grey bands mark the night period. The x-axis represents elapsed time (hh). Promoter region in the construct spanned -4.4kb to 0.2 kb relative to TSS.

Because drought alone elicited only a weak transcriptional response in this experiment (Suppl. Table S11), we restricted the memory analysis to the combined heat–drought treatment only. The transcriptional responses of first stress cycle correlated well with our initial experiment (Spearman rho = 0.56, 86% match in direction), and re-watering produced the expected switch from proline synthesis to proline breakdown (Yoshiba et al., 1997). We classified genes by their response to the first drought and its change in the second (Suppl. Table S9), following established classification scheme for transcriptional memory response-patterns (Ding et al., 2013). Genes repressed more strongly in the second drought ([-/-]) were enriched in photosynthesis, Calvin cycle, and starch biosynthesis, while genes induced more strongly ([+/+]) were enriched in vesicle trafficking and autophagy, Ca²⁺ signalling, ethylene action, and cell wall modification, among others (Suppl. Table S10). Genes repressed in the first drought whose response rose in the second ([-/+]) were enriched in NAC and WRKY transcription factors and pathogen response, including salicylic acid immunity regulators, and in further Ca²⁺ signalling components beyond those amplified in the [+/+] class. The ABA signalling and dehydration response, including *StABF2*, was induced to a similar extent in both cycles ([+/=]). While ABA signalling carried no memory response, ABA metabolism did: the rate-limiting biosynthetic enzyme *StNCED3* and the ABA 8’-hydroxylase *StCYP707A1* were classified as [+/+] indicating that ABA metabolism is primed for stronger response in the second stress occurrence.

Stronger re-induction has been attributed to chromatin that remains accessible after the first stress (Kappel et al., 2023), but has only recently been tested genome-wide using ATAC-seq (Z.-W. Wang et al., 2026). We compared the [+/+] genes with [+/=] genes, considering accessibility from our initial heat–drought experiment. On average, no greater changes in chromatin accessibility were observed around memory genes than around any matched genes in both cultivars. Nevertheless, memory genes contained significantly more accessible regions, located upstream of their TSS and in the gene bodies (Fig. 4B, Suppl. Fig. S10). *StNCED3* illustrates this pattern, with thirteen accessible regions spanning this locus and its TSS region ranking among the top 1.2 % most accessible regions across control and stress-treated samples. Because accessibility changes varied across regions, enhanced re-induction of *StNCED3* may depend primarily on the availability of transcriptional regulators rather than additional chromatin opening. To identify potential regulators, we performed a network analysis using the Comprehensive Knowledge Network (CKN) (Bleker et al., 2024). A curated list of potato genes involved in ABA biosynthesis (Suppl. Table S12) was used as seed nodes to identification of their interacting partners and construct a transcriptional regulatory subnetwork. This subnetwork was overlaid with transcriptomics data from the priming experiment showing that *StNCED3* was mildly upregulated in the first drought cycle, while *StWRKY33b* was not regulated (Fig. 4C). Regulation of both genes was enhanced in the second round of drought (Fig. 4C, left vs right panel) indicating StWRKY33b may contribute to priming of potato plants.

To test the hypothesis that StWRKY33b regulates StNCED3, we first examined chromatin accessibility at both loci. Accessibility of the *StWRKY33b* promoter increased slightly, but changes were not statistically significant (Suppl. Table S5). Three accessible regions were detected within the 4.5 kb *StNCED3* promoter, none of which changed significantly in either treatment; the strongest change at the locus was a decrease in accessibility 8.7 kb upstream (Fig. 4D, Suppl. Table S5). A binding motif for WRKY transcription factors was identified in the TSS-proximal accessible region (Fig. 4D). To test whether the enhanced induction of *StNCED3* upon recurring heat–drought stress could be driven by StWRKY33b, we performed transactivation studies in *Nicotiana benthamiana*. A strong activation of *StNCED3* gene expression by StWRKY33b was confirmed in these assays, which can explain the plant’s response to reoccurring stress (Fig. 4E, Suppl. Table S13).

## DISCUSSION

Responses of plants to combined and recurrent stresses cannot be predicted from single-stress responses, and their regulatory basis, including the contribution of chromatin accessibility, remains largely unknown, particularly in crops such as potato. Our integrated chromatin accessibility and transcriptome analysis reveals shared and stress-specific responses to heat, drought and waterlogging and their combination, and shows how a previous stress exposure primes ABA biosynthesis through StWRKY33b.

Approximately two thirds of the stress-responsive chromatin regions were specific to a single stress (Fig. 1D). Drought, heat–drought and waterlogging shared a substantial set of differentially accessible regions, with regions exhibiting increased accessibility under all three conditions associated with canonical ABA- and dehydration-responsive genes (Fig. 1D; Suppl. Table S5). This shared response signature is consistent with our previous physiology and hormone profiling studies of Désirée, where waterlogging activated ABA responses similar to those induced by drought (Zagorščak et al., 2025). Leaves of waterlogged plants experience water deficit, because oxygen deprivation reduces root water uptake through aquaporins closure (Tournaire-Roux et al., 2003). Our data extend these observations by showing that drought and waterlogging also share regulatory responses at the chromatin level. The overlap was nevertheless partial. The ABRE motif, recognized by group A bZIP transcription factors, gained accessibility genome-wide under drought and heat–drought, but only modestly under waterlogging, where WRKY and NAC motifs predominated (Fig. 2B; Suppl. Fig. S6). Waterlogging thus increases accessibility of subset of ABA-responsive genes, including *StABF2* itself, without triggering the widespread increase in ABRE accessibility seen under drought.

Heat had the weakest effect on chromatin accessibility in Désirée, and heat-treated Lady Rosetta plants were barely distinguishable from controls (Fig. 1C; Suppl. Fig. S3). We did not observe the canonical heat-shock signature reported in Arabidopsis, where acute heat shock alters accessibility at thousands of regulatory elements for up to two days (Sullivan et al., 2014). However, these differences may reflect the experimental conditions. Arabidopsis studies were done on young (7 day-old) seedlings exposed to severe heat shock, whereas our experiments were carried out on fully grown (4-week-old) plants subjected to prolonged, moderate heat stress. Our findings are consistent with the limited changes in DNA methylation reported in two other potato cultivars exposed to prolonged, albeit somewhat more severe, heat stress at a similar developmental stage (Yeo et al., 2026). Nevertheless, strong transcriptional induction of selected heat shock protein (*HSP*) genes was observed after prolonged heat stress exposure, and HSF binding motifs were identified in their regulatory regions (Fig. 2B, Suppl. Table S4).

Genome-wide changes in chromatin accessibility and gene expression under combined heat– drought stress more closely resembled those induced by drought than by heat, particularly in Lady Rosetta (Fig. 1E, Fig. 3A). Nonetheless, genes exclusively responding to combined heat–drought stress represented the largest response class (Fig. 3B). Thus, despite the stronger influence of drought, combined stress elicited a largely unique transcriptional response, in agreement with previous combined heat and drought stress studies in potato, tobacco and Arabidopsis (Rizhsky et al., 2002, 2004; Zagorščak et al., 2025).

The sequential stress treatment design may partly explain why the combined heat–drought response more closely resembled drought than heat. Plants were exposed to heat before drought was imposed, which might have already affected their response. Combined stress enhanced the expression of genes involved in protein quality control. While changes in chromatin accessibility were restricted to specific genes, transcriptional induction of chaperones, including *StHSP90,* roughly doubled under heat–drought compared to drought (Suppl. Fig. S7). This enhanced response may reflect higher leaf temperature: drought-induced stomatal closure limits transpiration cooling, causing leaves to become warmer than under heat alone (Rizhsky et al., 2002; Zagorščak et al., 2025). In contrast, the ABF group of transcriptional regulators, which act as key regulators downstream of ABA signalling, showed weaker transcriptional induction under heat–drought than under drought alone. Chromatin accessibility at these genes was also reduced in Désirée but not in Lady Rosetta (Fig. 3C). At the *StABF2* locus, this reduction under heat–drought occurred mainly at distal regions 1–5 kb upstream of the TSS, whereas accessibility at the proximal promoter region changed only slightly (Fig. 3D). Combined stress may therefore alter *StABF2* regulation through distal regulatory regions rather than through changes in the TSS-proximal promoter. Interestingly, most genes showing combined stress-specific changes in chromatin accessibility in both cultivars were linked to stress signalling, including Ca^2+^ signalling, proteasome-mediated protein degradation, and hormonal signalling pathways. Approximately one third of these genes, however, remain functionally uncharacterized (Suppl. Table S5).

Within an agricultural context, crops frequently experience multiple environmental stresses, either individually or in combination, with repeated cycles of stress and recovery throughout the growing season. We therefore designed a stress-priming experiment to identify novel regulators of heat and drought that provided a stress memory which may enhance responses to recurrent stress. Stronger transcriptional induction following a previous stress has been commonly attributed to sustained chromatin accessibility (Kappel et al., 2023), or the activity of specific transcriptional factors (Crawford et al., 2026; Jiang et al., 2025; Wei et al., 2026). Our data suggest that in potato leaves exposed to prolonged combined stress, pre-existing chromatin accessibility, rather than stress-induced chromatin opening, identifies genes exhibiting transcriptional memory. This is consistent with a recent study showing that stress-responsive regulatory elements are concentrated in constitutively accessible TSS-proximal regions (Morales-Cruz et al., 2026). Thus, pre-existing chromatin accessibility may provide a permissive regulatory state facilitating transcriptional memory upon subsequent stress exposure.

*StNCED3,* which encodes an enzyme catalysing the rate-limiting step of ABA synthesis (Vo et al., 2025), supports this model. Its promoter remains constitutively accessible, suggesting that its enhanced re-induction depends primarily on its transcriptional regulators rather than changes in chromatin. By integrating a regulatory knowledge network with the gene expression data, we identified AtWRKY33 as a key regulator of *AtNCED3* (Fig. 4C). AtWRKY33 was shown to transcriptionally repress *AtNCED3* in *Arabidopsis* (S. Liu et al., 2015). In potato, we identified two *AtWRKY33* orthologs, *StWRKY33a* and *StWRKY33b* (Suppl. Fig. S11), while only *StWRKY33b* exhibited enhanced transcriptional induction upon recurrent heat–drought stress in our study (Fig. 4C). Our transactivation assays further demonstrated that StWRKY33b directly activates *StNCED3* transcription (Fig. 4E), thereby placing stress memory on the upstream regulators of *StNCED3*. The activating rather than repressive function of StWRKY33b may reflect a functional divergence from AtWRKY33 or a context-dependent switch in its repressor/activator role, as already demonstrated for AtWRKY33 (S. Liu et al., 2015).

StWRKY33b regulation of heat and drought responses is particularly relevant for a heat-sensitive crop such as potato (Guillemette et al., 2024), as concurrent heat and drought events are becoming more frequent and severe. Our findings show that prolonged abiotic stress leaves a lasting, largely stress-specific imprint on the chromatin landscape of potato, and that the imprint of combined heat and drought stresses is very different compared to those of single stresses. This imprint allows appropriate response to prolonged stress and encodes a memory of past exposure that primes plants for stress reoccurrence. As heat and drought cycles become increasingly more common within a single growing season, such stress memory may become a decisive determinant of future potato crop performance. Resolving how it is established and maintained will open new routes to breeding crops that are resilient not only to individual stresses but to the compound and reoccurring stress regimes of a changing climate.

## METHODS

### Plant growth

Plant growth and stress application followed the protocol detailed in Abdelhakim et al. (Abdelhakim et al., 2024). Potato tissue culture plantlets (cvs. Désirée and Lady Rosetta) were propagated in 90 mm petri dishes containing MS medium (Murashige & Skoog, 1962) supplemented with 20 g/l sucrose and 8 g/l agar at 18 °C ± 4 °C, 16 h light/ 8 h dark, and light intensity 100 µmol m^-^² s^-^¹. Four weeks after sub-culture, *in vitro* plantlets were transferred to 250 ml pots containing Klasmann Substrate 2 and kept in a climate-controlled chamber under low light for 1 week. After 10 days, plants were transplanted into 3L pots containing 3:1 Klasmann Substrate 2: sand and grown in a growth chamber at 320 µmol m⁻² s⁻¹ (16 h photoperiod) and 22/19°C day/night. Stress treatments were imposed at the early tuberisation stage, 28 days after transplanting the in vitro cuttings. Relative humidity was maintained at 55% throughout the experiment, including in the heat-treatment chamber.

For the initial abiotic stress experiment, plants were exposed to three individual stresses: heat (30/28 °C), drought (20% soil relative water content, SRWC), and waterlogging (160% SRWC), and a combined heat–drought treatment. Control plants were maintained at 22/19 °C and 60% SRWC.

For the priming experiment, plants were maintained either at 18 or 28 °C. At each temperature, plants were subjected to 4 days of drought or were watered once daily to maintain the field capacity (Suppl. methods and results, Suppl. Fig. S8). After four days of treatment, all plants were sampled, and the drought-treated plants were then watered to field capacity for one day to allow recovery. Plants were sampled again after one day of recovery and then either subjected to a second period of 4 days of drought or watered daily to field capacity prior to the final harvest. Details on sample treatment groups and experimental design are given in Suppl. Table S6. Soil moisture, relative water content, and photosynthetic and stomatal parameters were measured as described in Suppl. methods and results.

### ATAC-seq sampling and library preparation

At harvest, the second leaf of each plant was collected. Nuclei were isolated straight away from fresh tissue. Plants were processed in batches of four, with treatment groups alternated within each batch. For each sample, approximately 150 mg of leaf tissue with the midvein removed was chopped on ice for 1.5 min (200–250 strokes min⁻¹) with a fresh double-edged razor blade. Chopping was done in 500 µl of ice-cold extraction buffer: 0.25 M sucrose, 10 mM Tris-HCl pH 8.0, 10 mM MgCl₂, 0.5% Triton X-100, 5 mM β-mercaptoethanol and protease inhibitor (Roche). The homogenate was rinsed with 500 µl of buffer and filtered through 100 µm and then 40 µm cell strainers (pluriStrainer Mini, pluriSelect). A low-speed spin (50 × g, 5 min, 4 °C) removed large debris. Nuclei were then pelleted at 300 × g for 5 min at 4 °C and washed twice in 1 ml of extraction buffer. Tagmentation and library preparation followed the instructions for the ATAC-seq Kit (Diagenode). Briefly, washed nuclei were resuspended in 50 µl of tagmentation mix containing 2.5 µl of loaded Tagmentase (Tn5), 0.01% digitonin and 0.1% Tween-20, then incubated for 30 min at 37 °C. The reaction was stopped with 300 µl of Buffer ERC. Tagmented DNA was purified with the MinElute Reaction Cleanup Kit (Qiagen) and eluted in 10 µl of Buffer EB.

In the preamplification step, tagmented DNA was amplified by PCR with indexed primers (Diagenode) and Phusion High-Fidelity DNA Polymerase (NEB), with an extension time of 30 s. To determine the number of additional amplification cycles required, qPCR was performed with an aliquot of first-round PCR product, Phusion High-Fidelity DNA Polymerase (NEB) and 1× SYBR Green (Invitrogen). The quantification cycle value corresponding to one-third of the maximum fluorescence was used to determine the number of additional PCR cycles applied to pre-amplified libraries. The pre-amplified libraries were then subjected to the additional PCR cycles. Product size distributions were assessed on a TapeStation system with the D5000 ScreenTape Assay (Agilent Technologies). Final PCR products were purified with 1.8x AMPure XP beads (Beckman Coulter). DNA concentrations were measured on a Qubit fluorometer with the Qubit dsDNA High Sensitivity Assay Kit (Thermo Fisher Scientific).

### ATAC-seq data analysis

Raw reads were quality-checked with FastQC v0.11.9 (Andrews, 2010) and trimmed for Tn5 mosaic-end adapters using cutadapt v4.8 (Martin, 2011) in paired-end mode. Reads were aligned to the DM v6.1 genome carrying the UniTato gene models (Zagorščak et al., 2024), supplemented with the complete Désirée organellar genome, comprising the plastid (NCBI DQ386163.2) and the three molecules of the multipartite mitochondrial genome (NCBI MN104801.1, MN104802.1 and MN104803.1), using BWA-MEM v0.7.17-r1188 (H. Li, 2013).

Alignments were sorted and indexed with SAMtools v1.20 (Danecek et al., 2021), and PCR duplicates were flagged with samblaster v0.1.26 (Faust & Hall, 2014). Organellar reads were removed before downstream analysis. Unless otherwise stated, analyses used reads with MAPQ ≥ 40 and excluded duplicate reads.

Dataset quality was assessed using SAMtools stats and deepTools v3.5.4 (Ramírez et al., 2016), including fingerprinting, sample correlation and principal component analysis, fraction of reads in peaks (FRiP), and TSS enrichment. Coverage tracks were generated using alignmentSieve (deepTools v3.5.4) with ATAC-seq strand shifts of +4/−5 bp, followed by bamCoverage (deepTools v3.5.4) with 1-bp bins and RPGC normalisation using an effective genome size of 600 Mb. FRiP was calculated relative to the consensus peak set.

Two libraries were excluded: LR-0501-Dc3, which yielded too few usable reads (0.16 million reads in consensus regions, against at least 1.45 million in every other library; duplicate rate 82.1%; TSS enrichment 2.0), and De-0620-C4, whose library required 20 PCR amplification cycles, against 12 or fewer for every other library. The final dataset comprised 54 of 56 libraries. Peaks were called separately for each cultivar and treatment group using Genrich v0.6.1 (Gaspar, 2018) in paired-end mode with MAPQ ≥40, duplicate removal, and q ≤0.05. All libraries except LR-0501-Dc3 were used for peak calling. A fixed-width consensus peak universe was generated as non-overlapping 300-bp windows centred on Genrich summits, retaining the higher-signal summit where windows overlapped. The window size was based on the observed peak-width distributions (Suppl. Fig. S2A). Before the consensus was built, peaks whose read counts across libraries tracked the library organellar read fraction (92 to 250 peaks per treatment group), indicating plastid-derived reads mapped to nuclear plastid-like insertions, were removed.

Reads were quantified with featureCounts v2.0.6 (Liao et al., 2014) in paired-end mode using a minimum fractional overlap of 0.2. Regions were annotated against the UniTato gene models using ChIPseeker v1.48.0 (Yu et al., 2015) with a ±3-kb promoter window. Differential accessibility analysis was performed in R v4.6.0 using edgeR v4.10.1 (Robinson et al., 2010) *and limma v3.68.4* (Ritchie et al., 2015). Regions were filtered with filterByExpr (min.count = 50). GC-content bias was removed with conditional quantile normalisation (cqn) (Hansen et al., 2012), and the resulting offsets were supplied to voomLmFit (Law et al., 2014) using a one-way design, followed by empirical Bayes moderation (eBayes) with robust estimation and Benjamini–Hochberg correction (Benjamini & Hochberg, 1995). Differentially accessible regions were defined as FDR <0.05 and |log2 fold change| >0.5. Per-cultivar, cultivar-averaged, stress-by-cultivar, and direct combined-stress versus drought contrasts were extracted from the same model.

Sample structure was assessed by multidimensional scaling of the 5,000 most variable regions. Response magnitude was summarised from the distribution of absolute log2 fold changes across tested regions, and similarity between contrasts was assessed by Pearson correlation of log_2_ fold changes.

For baseline accessibility analyses, a gene was considered to have an accessible promoter if a consensus region lay within 1 kb or 3 kb of its TSS in at least one of the three control harvest groups of that cultivar. Proportions were computed over all 57,394 annotated genes and over the 22,134 genes retained by the expression filter of the RNA-seq analysis.

### RNA extraction and sequencing

For the initial experiment RNA-seq analysis, the 3^rd^ leaf of same plants as for the ATAC-seq analysis was sampled. Leaf tissue was homogenised under liquid nitrogen using a mortar and pestle, and total RNA was extracted with MagMAX Plant RNA Isolation Kit (Thermo Fisher Scientific) on a KingFisher Apex purification system (Thermo Fisher Scientific), including DNase I treatment (Thermo Fisher Scientific). Reverse-stranded, polyA-enriched Illumina RNA-seq libraries were prepared and sequenced in paired-end, 150 bp mode on an Illumina NovaSeq X Plus (Novogene). Read adapter trimming was performed by the sequencing service prior to data delivery.

For the priming experiment RNA-seq analysis, fully expanded leaves were harvested from the fourth node of three independent replicate plants and immediately frozen in liquid nitrogen. Samples were stored at −80 °C prior to RNA extraction. Total RNA was extracted from potato leaves using RNeasy Plant Mini Kit (Qiagen) as described previously (Ducreux et al., 2008). Reverse-stranded, polyA-enriched Illumina RNA-seq libraries were prepared and sequenced in single-end, 100 bp mode on an Illumina HiSeq 2500 by IGA Technology Services (Udine, Italy).

### Differential expression analysis

For transcript quantification from both RNA-seq experiments, a decoy-aware index using the potato *Solanum tuberosum* Group Phureja DM genome (Pham et al., 2020) and the UniTato transcriptome (Zagorščak et al., 2024) was built and reads were quantified with Salmon v1.10.3 (Patro et al., 2017). Mapping statistics were aggregated with MultiQC (Ewels et al., 2016). Two samples from the initial experiment RNA-seq with outlying extremely low mapping rates were discarded, leaving 246 out of total 248 samples. For the priming experiment, prior to pseudomapping, adapter removal was performed using Trim Galore v2.2.0 given the parameters “--quality 20 --polyA --trim-n”.

Differential expression analysis was performed in R v4.6.0 using edgeR v4.10.1 (Chen et al., 2025). Briefly, transcript-level abundances were imported and summarised to gene level counts with tximport v1.40.0 (Soneson et al., 2016), lowly expressed genes were removed with *filterByExpr* (min.count = 50, min.total.count = 100), counts were TMM-normalised and modelled with *voomLmFit* (sample.weights = TRUE; Baldoni et al., 2025), and predefined contrasts were tested with *eBayes* using limma v3.68.4 (Ritchie et al., 2015). Differentially expressed genes were defined as Benjamini–Hochberg-adjusted P <0.05 and, where applied, |log_2_ fold change| >1. For the priming experiment, drought-memory classes were assigned to genes in the 28 °C treatment following an established scheme (Ding et al., 2013). Each gene was classified as [HD1/HD2 − HD1], where HD1 is the response to the first drought and HD2 – HD1 is the fitted difference between the second and the first drought response, both relative to the well-watered 28 °C control sampled at the same time point. Each term was scored as significantly positive (+), significantly negative (-) or not significant (=) at FDR < 0.05.

### Integration of RNA-seq and ATAC-seq

Accessibility and expression data were paired by cultivar, stress, and sampling day. Genes were linked to ATAC-seq regions using the ChIPseeker annotation, and one representative promoter region was retained per gene, selected by the lowest accessibility FDR and, for ties, the largest absolute log_2_ fold change.

For genes that were differentially expressed and differentially accessible in both assays (FDR < 0.05), directional concordance was quantified as the proportion showing changes in the same direction. Coupling between accessibility and expression changes was assessed across all genes with a tested promoter peak and an RNA-seq result, without selection on significance or direction, as the Pearson correlation (with 95% confidence interval) and the Spearman rank correlation of the log2 fold changes, separately for each cultivar and stress. The overlap between differentially expressed genes and genes with a differentially accessible promoter was counted over gene-by-comparison tests, pooled across the four stresses and both cultivars.

Combined-stress responses were compared with each constituent individual stress, separately for transcripts (per gene) and for chromatin accessibility (per accessible region). A gene or region was considered responsive at FDR < 0.05 and |log_2_FC| > 1 for transcripts or |log_2_FC| > 0.5 for accessibility. Genes or regions responsive to either treatment were classified as emergent when responsive only under combined stress, blunted when responsive only under the individual stress, redirected when the direction of change differed, amplified or attenuated when changes were in the same direction but the combined-to-individual log2 fold-change ratio was >1.2 or <0.8, respectively, and retained otherwise.

### Gene set enrichment analyses

Gene set enrichment analysis was performed using UniTato Mercator4 v7 (MapMan4) functional bins (Zagorščak et al., 2024). For ATAC-seq, normalised counts of a single promoter-associated region per gene with the highest mean consensus Genrich q-value was used. For RNA-seq, gene-level normalised counts per million were used as input.

Enrichment was calculated using an R implementation of the Broad’s GSEA (Subramanian et al., 2005) algorithm with signal-to-noise ranking, 1,000 gene-label permutations, gene-set sizes of 15–500, and the unweighted Kolmogorov–Smirnov statistic. A fixed random seed was used, and FDR q-values were adjusted separately for each direction of enrichment. Gene sets with FDR <0.05 were considered enriched; FDR 0.05–0.10 was considered suggestive. For display (Fig. 2A, Suppl. Fig. S5), bins with FDR <0.10 in the accessibility data in at least two of the eight comparisons were selected, and nested bins were reduced to the most general qualifying bin.

Because the ATAC-seq analysis was restricted to genes with annotated promoter regions, the accessibility and expression analyses used different gene universes. Enrichment results were therefore compared by direction rather than magnitude.

To compare drought and combined-stress responses, genes were ranked using the signed statistic

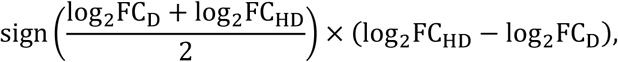

where D and HD denote drought and combined heat and drought responses, respectively. MapMan bins were tested using fgsea (Korotkevich et al., 2019) with gene-set sizes of 5–500 and 10,000 permutations. The gene universe comprised the union of drought- and combined-stress-responsive genes, oriented according to their mean response. The same procedure and gene sets were applied to the accessibility data using the fitted combined-stress versus drought contrast at promoter-associated regions.

Over-representation of MapMan4 bins (levels 2 and 3, 15–600 genes per bin) in the [+/+], [+/-], [-/-] and [-/+] drought-memory classes was tested against all genes changed in the same direction in HD1, using a Cochran–Mantel–Haenszel test stratified by quartiles of absolute HD1 log_2_FC, so that each class is compared with genes of similar first-drought response. Bins with at least five genes in the class were tested, and FDR was controlled with the Benjamini– Hochberg procedure within each class.

### Motif enrichment

Motif sequences from JASPAR CORE Plants 2026 (Ovek Baydar et al., 2026) were scanned against consensus peak sequences using FIMO (MEME Suite v5.0.5; Grant et al., 2011) with P ≤1 × 10⁻⁴. The FIMO hits served as the motif annotation for chromVAR and mark the motif positions in Fig. 4D. Redundant JASPAR motif models were grouped into motif archetypes by cutting the precomputed RSAT matrix-clustering hierarchy of the JASPAR 2026 plants CORE collection (Castro-Mondragon et al., 2017) at a normalised correlation (Ncor) of 0.70.

Differential motif activity was assessed with chromVAR v1.34.1 (Schep et al., 2017) using per-region GC content as a bias term and 2,000 background iterations, each drawing a GC- and accessibility-matched background region for every peak. Bias-corrected deviation z-scores were quantile-normalised, and differential activity for stress-versus-control comparisons was tested with limma v3.68.4 (Ritchie et al., 2015). A linear model was fitted separately for each cultivar, stress-versus-control contrasts were extracted from it, and standard errors were moderated with eBayes. P-values were Benjamini–Hochberg adjusted across motifs within each cultivar and contrast. Motifs occurring in fewer than 10 regions were excluded.

### Network-based analyses

A list of 72 potato genes involved in isoprenoid biosynthesis (seed genes) was compiled, and their identifiers were unified to DM v6.1 gene identifiers using the UniTato translation table (Zagorščak et al., 2024). Their Arabidopsis orthologues were then identified using PLAZA 5.0 (Van Bel et al., 2022), OrthoDB (Kuznetsov et al., 2023), and reciprocal BLAST hits (Benoit & Drost, 2021). Pairs with inconsistent MapMan4 (Schwacke et al., 2019) annotation were excluded, leaving 48 Arabidopsis seed genes.

Regulatory interactions of ranks 0–2 were taken from AtCKN v2 (Bleker et al., 2024). The first-order neighbourhood of the seed genes was extracted in R v4.5.2 (R Core Team, 2026) using igraph v2.3.3 (Antonov et al., 2023), and the largest weakly connected component was retained. The network was restricted to transcription factor regulation, retaining edges from transcription factors to seed genes and edges between these transcription factors. Using the priming experiment RNA-seq dataset, each node was mapped to its most strongly differentially expressed potato orthologue (highest absolute log_2_FC in the majority of comparisons) and annotated with its expression changes. The network was visualised in Cytoscape v3.9.1(Shannon et al., 2003).

### Multiple sequence alignment analysis

To classify the identified novel regulators of ABA synthesis, we performed additional multiple sequence alignment analysis. The orthology tables prepared for the network analyses returned many-to-many mappings between AtWRKY33, its close paralogues (AtWRKY25 and AtWRKY26) and multiple Solanaceae WRKYs. The potato and tomato orthologues of AtWRKY33 were therefore determined by phylogenetic analysis of group I WRKY proteins, which are defined by two WRKY domains, each consisting of a W[R/K]KYG[Q/K/E]K motif followed by an intact C2H2 zinc finger (Eulgem et al., 2000). Group I WRKY proteins were selected from *Arabidopsis thaliana* Araport11 (Cheng et al., 2017), *Solanum tuberosum* UniTato (Zagorščak et al., 2024) and *Solanum lycopersicum* ITAG5.0 (ITAG5.0; Y. Zhou et al., 2022) annotations, using the longest isoform of each gene. Pairwise global alignments of the full-length proteins were computed with pwalign v1.6.0 (Aboyoun & Gentleman, 2025) (BLOSUM62, gap opening 10, gap extension 0.5), and distances were calculated as 1 − sequence identity. A neighbour-joining tree was built with ape v5.8-1 (Paradis & Schliep, 2019), rooted on AtWRKY54 (group III), and divided into 10 clusters, equal to the number of Arabidopsis sequences including the outgroup, by repeatedly splitting the deepest clade starting from the root (Suppl. Fig. S11). AtWRKY33 clustered with two genes from each Solanaceae species, corresponding in tomato to SlWRKY33a and SlWRKY33b (J. Zhou et al., 2015). The potato orthologues were named StWRKY33a (Soltu.DM.09G009490) and StWRKY33b (Soltu.DM.06G018840). Analyses were performed in R v4.5.2, and the phylogenetic tree was drawn with ggtree v4.0.5 (Xu et al., 2022).

### Transient dual-luciferase reporter system

The putative promoter region of the nine-cis-epoxycarotenoid dioxygenase 3 (*StNCED3*; Soltu.DM.07G022620) gene from the *Solanum tuberosum* group Phureja DM1-3 v6.1 genome was amplified utilizing forward primer: TGGTAGTGCAAATTTTATGTTATGTC and reverse primer: TTTGTAGCATGTGAAGTAGTAGTGGC targeting 4.5 kb upstream of the TSS. The resulting amplified promoter product was then cloned into the GATEWAY cloning vector pDONR207, which was subsequently inserted into the pGWB435 destination vector. The pGWB435 vector harboured a luciferase reporter gene, with incorporation of the potato intron of the L700 gene using the LR clonase reaction (Invitrogen) to complete the construct.

Reporter construct *35S::StWRKY33b* was cloned and transformed into *Agrobacterium tumefaciens* strain GV3101, alongside the *pStNCED3::iLUC* construct. The construct was co-infiltrated into *Nicotiana benthamiana* (rdr6i) leaves. The Agrobacterium strain carrying the *pStNCED3-iLUC* fusion in conjunction with the *35S::StWRKY33b* construct was suspended in a solution containing 10 mM MES (pH 5.8), 10 mM MgSO4, and 150 µM acetosyringone, with an optical density of 0.2 at 600 nm for all cultures utilized in infiltration.

Two days post-infiltration, leaf discs were incubated in a plant medium composed of MS+2% sucrose, supplemented with 0.02 mg/mL D-luciferin (Promega). Luciferase activity was monitored every 30 minutes using a microplate luminometer Centro LB-960 (Berthold Technologies), and the data were captured using MikroWin 2000 v4.29 software (Mikrotek Laborsysteme).

## ACKNOWLEDGEMENTS

We acknowledge Luka Kranjc and Sandi Hunyadi for the help with data management and deposition.

## AUTHOR CONTRIBUTIONS

S.S., R.D.H., M.A.T., S.P., M.T. and K.G. conceived the study. All authors contributed to the development of the methodology. T.G., L.A., R.C. and M.A.S. performed the experiments. K.P., M.A.T., S.P. and K.G. provided resources. T.G., M.Z., Ž.R. and M.P. developed the computational pipelines. T.G., M.Z., M.P. and K.G. performed the formal analysis and curated the data. T.G., M.Z. and K.G. prepared the figures. T.G., M.Z., M.P. and K.G. wrote the original draft of the manuscript, and all authors reviewed and edited it. R.D.H., M.A.T., S.P., M.T. and K.G. supervised the research, administered the project and acquired funding.

## DATA AVAILABILITY

Raw reads from the initial and priming experiment were deposited in the European Nucleotide Archive (ENA) at EMBL-EBI under accession numbers PRJEB127277 and PRJEB129741, respectively.

## REFERENCES

Abdelhakim, L. O. A., Pleskačová, B., Rodriguez-Granados, N. Y., Sasidharan, R., Perez-Borroto, L. S., Sonnewald, S., Gruden, K., Vothknecht, U. C., Teige, M., & Panzarová, K. (2024). High Throughput Image-Based Phenotyping for Determining Morphological and Physiological Responses to Single and Combined Stresses in Potato. Journal of Visualized Experiments, (208), 66255. 10.3791/66255

Aboyoun, P., & Gentleman, R. (2025). pwalign: Perform pairwise sequence alignments (Version 1.6.0) [Computer software]. Bioconductor. 10.18129/B9.bioc.pwalign

Andrews, S. (2010). S-andrews/FastQC: A quality control analysis tool for high throughput sequencing data. https://github.com/s-andrews/FastQC

Antonov, M., Csárdi, G., Horvát, S., Müller, K., Nepusz, T., Noom, D., Salmon, M., Traag, V., Welles, B. F., & Zanini, F. (2023). *Igraph enables fast and robust network analysis across programming languages* (Version 1). arXiv. 10.48550/ARXIV.2311.10260

Baldoni, P. L., Chen, L., Li, M., Chen, Y., & Smyth, G. K. (2025). Dividing out quantification uncertainty enables assessment of differential transcript usage with limma and edgeR. Nucleic Acids Research, 53(22), gkaf1305. 10.1093/nar/gkaf1305

Benjamini, Y., & Hochberg, Y. (1995). Controlling the False Discovery Rate: A Practical and Powerful Approach to Multiple Testing. Journal of the Royal Statistical Society Series B: Statistical Methodology, 57(1), 289–300. 10.1111/j.2517-6161.1995.tb02031.x

Benoit, M., & Drost, H.-G. (2021). A Predictive Approach to Infer the Activity and Natural Variation of Retrotransposon Families in Plants. In J. Cho (Ed.), Plant Transposable Elements (Vol. 2250, pp. 1–14). Springer US. 10.1007/978-1-0716-1134-0_1

Bleker, C., Ramšak, Ž., Bittner, A., Podpečan, V., Zagorščak, M., Wurzinger, B., Baebler, Š., Petek, M., Križnik, M., Van Dieren, A., Gruber, J., Afjehi-Sadat, L., Weckwerth, W., Županič, A., Teige, M., Vothknecht, U. C., & Gruden, K. (2024). Stress Knowledge Map: A knowledge graph resource for systems biology analysis of plant stress responses. Plant Communications, 5(6), 100920. 10.1016/j.xplc.2024.100920

Buenrostro, J. D., Giresi, P. G., Zaba, L. C., Chang, H. Y., & Greenleaf, W. J. (2013). Transposition of native chromatin for fast and sensitive epigenomic profiling of open chromatin, DNA-binding proteins and nucleosome position. Nature Methods, 10(12), Article 12. 10.1038/nmeth.2688

Castro-Mondragon, J. A., Jaeger, S., Thieffry, D., Thomas-Chollier, M., & van Helden, J. (2017). RSAT matrix-clustering: Dynamic exploration and redundancy reduction of transcription factor binding motif collections. Nucleic Acids Research, 45(13), e119– e119. 10.1093/nar/gkx314

Chen, Y., Chen, L., Lun, A. T. L., Baldoni, P. L., & Smyth, G. K. (2025). edgeR v4: Powerful differential analysis of sequencing data with expanded functionality and improved support for small counts and larger datasets. Nucleic Acids Research, 53(2), gkaf018. 10.1093/nar/gkaf018

Cheng, C., Krishnakumar, V., Chan, A. P., Thibaud-Nissen, F., Schobel, S., & Town, C. D. (2017). Araport11: A complete reannotation of the *Arabidopsis thaliana* reference genome. The Plant Journal, 89(4), 789–804. 10.1111/tpj.13415

Crawford, T., Pratx, L., & Bäurle, I. (2026). Principles and mechanisms of plant acclimation to heat stress. Nature Reviews Molecular Cell Biology, 27(9), 668–683. 10.1038/s41580-026-00977-x

Danecek, P., Bonfield, J. K., Liddle, J., Marshall, J., Ohan, V., Pollard, M. O., Whitwham, A., Keane, T., McCarthy, S. A., Davies, R. M., & Li, H. (2021). Twelve years of SAMtools and BCFtools. GigaScience, 10(2), giab008. 10.1093/gigascience/giab008

Devaux, A., Goffart, J.-P., Petsakos, A., Kromann, P., Gatto, M., Okello, J., Suarez, V., & Hareau, G. (2020). Global Food Security, Contributions from Sustainable Potato Agri-Food Systems. In H. Campos & O. Ortiz (Eds.), The Potato Crop: Its Agricultural, Nutritional and Social Contribution to Humankind (pp. 3–35). Springer International Publishing. 10.1007/978-3-030-28683-5_1

Ding, Y., Liu, N., Virlouvet, L., Riethoven, J.-J., Fromm, M., & Avramova, Z. (2013). Four distinct types of dehydration stress memory genes in Arabidopsis thaliana. BMC Plant Biology, 13(1), 229. 10.1186/1471-2229-13-229

Ducreux, L. J. M., Morris, W. L., Prosser, I. M., Morris, J. A., Beale, M. H., Wright, F., Shepherd, T., Bryan, G. J., Hedley, P. E., & Taylor, M. A. (2008). Expression profiling of potato germplasm differentiated in quality traits leads to the identification of candidate flavour and texture genes. Journal of Experimental Botany, 59(15), 4219– 4231. 10.1093/jxb/ern264

Eulgem, T., Rushton, P. J., Robatzek, S., & Somssich, I. E. (2000). The WRKY superfamily of plant transcription factors. Trends in Plant Science, 5(5), 199–206. 10.1016/S1360-1385(00)01600-9

Ewels, P., Magnusson, M., Lundin, S., & Käller, M. (2016). MultiQC: summarize analysis results for multiple tools and samples in a single report. Bioinformatics, 32(19), 3047– 3048. 10.1093/bioinformatics/btw354

Faust, G. G., & Hall, I. M. (2014). *SAMBLASTER*: Fast duplicate marking and structural variant read extraction. Bioinformatics, 30(17), 2503–2505. 10.1093/bioinformatics/btu314

Gallusci, P., Agius, D. R., Moschou, P. N., Dobránszki, J., Kaiserli, E., & Martinelli, F. (2023). Deep inside the epigenetic memories of stressed plants. Trends in Plant Science, 28(2), 142–153. 10.1016/j.tplants.2022.09.004

Gaspar, J. M. (2018). jsh58/Genrich: Detecting sites of genomic enrichment. https://github.com/jsh58/Genrich

Grant, C. E., Bailey, T. L., & Noble, W. S. (2011). FIMO: scanning for occurrences of a given motif. Bioinformatics, 27(7), 1017–1018. 10.1093/bioinformatics/btr064

Guillemette, A. M., Hernández Casanova, G., Hamilton, J. P., Pokorná, E., Dobrev, P. I., Motyka, V., Rashotte, A. M., & Leisner, C. P. (2024). The physiological and molecular responses of potato tuberization to projected future elevated temperatures. Plant Physiology, 197(1), kiae664. 10.1093/plphys/kiae664

Guo, M., Yang, F., Zhu, L., Wang, L., Li, Z., Qi, Z., Fotopoulos, V., Yu, J., & Zhou, J. (2024). Loss of cold tolerance is conferred by absence of the WRKY34 promoter fragment during tomato evolution. Nature Communications, 15(1), 6667. 10.1038/s41467-024-51036-y

Hansen, K. D., Irizarry, R. A., & Wu, Z. (2012). Removing technical variability in RNA-seq data using conditional quantile normalization. Biostatistics, 13(2), 204–216. 10.1093/biostatistics/kxr054

Hsu, F.-C., Chou, M.-Y., Chou, S.-J., Li, Y.-R., Peng, H.-P., & Shih, M.-C. (2013). Submergence Confers Immunity Mediated by the WRKY22 Transcription Factor in Arabidopsis. The Plant Cell, 25(7), 2699–2713. 10.1105/tpc.113.114447

Jiang, Z., van Zanten, M., & Sasidharan, R. (2025). Mechanisms of plant acclimation to multiple abiotic stresses. Communications Biology, 8(1), 655. 10.1038/s42003-025-08077-w

Jiang, Z., Verhoeven, A., Li, Y., Geertsma, R., Sasidharan, R., & Van Zanten, M. (2026). Deciphering acclimation to sublethal combined and sequential abiotic stresses in *Arabidopsis thaliana*. Plant Physiology, 200(3), kiae581. 10.1093/plphys/kiae581

Kappel, C., Friedrich, T., Oberkofler, V., Jiang, L., Crawford, T., Lenhard, M., & Bäurle, I. (2023). Genomic and epigenomic determinants of heat stress-induced transcriptional memory in Arabidopsis. Genome Biology, 24(1), 129. 10.1186/s13059-023-02970-5

Korotkevich, G., Sukhov, V., & Sergushichev, A. (2019). Fast gene set enrichment analysis (p. 060012). bioRxiv. 10.1101/060012

Kurtz, J., Andino, R., Boraschi, D., Contreras-Garduño, J., Kachroo, A., Khan, I., Lanz Mendoza, H., Mukherjee, K., Peuß, R., & Ton, J. (2025). Trained immunity and immune priming in plants and invertebrates. eLife, 14, e106597. 10.7554/eLife.106597

Kuznetsov, D., Tegenfeldt, F., Manni, M., Seppey, M., Berkeley, M., Kriventseva, E. V., & Zdobnov, E. M. (2023). OrthoDB v11: Annotation of orthologs in the widest sampling of organismal diversity. Nucleic Acids Research, 51(D1), D445–D451. 10.1093/nar/gkac998

Law, C. W., Chen, Y., Shi, W., & Smyth, G. K. (2014). voom: Precision weights unlock linear model analysis tools for RNA-seq read counts. Genome Biology, 15(2), R29. 10.1186/gb-2014-15 2-r29

Li, H. (2013). Aligning sequence reads, clone sequences and assembly contigs with BWA-MEM (Version 2). arXiv. 10.48550/ARXIV.1303.3997

Li, X., Chen, L., Liu, T., Chen, Y., Wang, J., & Song, B. (2024). Integrated analysis of ATAC-seq and transcriptomic reveals the ScDof3*-ScproC* molecular module regulating the cold acclimation capacity of potato. Plant Physiology and Biochemistry, 210, 108576. 10.1016/j.plaphy.2024.108576

Liang, Z., Zhang, Q., Ji, C., Hu, G., Zhang, P., Wang, Y., Yang, L., & Gu, X. (2021). Reorganization of the 3D chromatin architecture of rice genomes during heat stress. BMC Biology, 19(1), 53. 10.1186/s12915-021-00996-4

Liao, Y., Smyth, G. K., & Shi, W. (2014). featureCounts: An efficient general purpose program for assigning sequence reads to genomic features. Bioinformatics, 30(7), 923–930. 10.1093/bioinformatics/btt656

Liu, H., Able, A. J., & Able, J. A. (2022). Priming crops for the future: Rewiring stress memory. Trends in Plant Science, 27(7), 699–716. 10.1016/j.tplants.2021.11.015

Liu, S., Kracher, B., Ziegler, J., Birkenbihl, R. P., & Somssich, I. E. (2015). Negative regulation of ABA signaling by WRKY33 is critical for Arabidopsis immunity towards Botrytis cinerea 2100. eLife, 4, e07295. 10.7554/eLife.07295

Lu, Z., Marand, A. P., Ricci, W. A., Ethridge, C. L., Zhang, X., & Schmitz, R. J. (2019). The prevalence, evolution and chromatin signatures of plant regulatory elements. Nature Plants, 5(12), Article 12. 10.1038/s41477-019-0548-z

Martin, M. (2011). Cutadapt removes adapter sequences from high-throughput sequencing reads. EMBnet.Journal, 17(1), 10. 10.14806/ej.17.1.200

Morales, A., De Boer, H. J., Douma, J. C., Elsen, S., Engels, S., Glimmerveen, T., Sajeev, N., Huber, M., Luimes, M., Luitjens, E., Raatjes, K., Hsieh, C., Teapal, J., Wildenbeest, T., Jiang, Z., Pareek, A., Singla-Pareek, S., Yin, X., Evers, J., … Sasidharan, R. (2022). Effects of sublethal single, simultaneous and sequential abiotic stresses on phenotypic traits of *Arabidopsis thaliana*. AoB PLANTS, 14(4), plac029. 10.1093/aobpla/plac029

Morales-Cruz, A., Greenblum, S. I., Wang, P., Zhang, Y., Yang, L., Daum, C. G., Johnson, J., Baumgart, L. A., & O’Malley, R. C. (2026). Positional grammar of transcription factor binding partitions developmental and stress-response regulation in plants. Nature Communications, 17(1), 9403. 10.1038/s41467-026-75947-0

Murashige, T., & Skoog, F. (1962). A Revised Medium for Rapid Growth and Bio Assays with Tobacco Tissue Cultures. Physiologia Plantarum, 15(3), 473–497. 10.1111/j.1399-3054.1962.tb08052.x

Ovek Baydar, D., Rauluseviciute, I., Aronsen, D. R., Blanc-Mathieu, R., Bonthuis, I., de Beukelaer, H., Ferenc, K., Jegou, A., Kumar, V., Lemma, R. B., Lucas, J., Pochon, M., Yun, C. M., Ramalingam, V., Deshpande, S. S., Patel, A., Marinov, G. K., Wang, A. T., Aguirre, A., … Mathelier, A. (2026). JASPAR 2026: Expansion of transcription factor binding profiles and integration of deep learning models. Nucleic Acids Research, 54(D1), D184–D193. 10.1093/nar/gkaf1209

Paradis, E., & Schliep, K. (2019). ape 5.0: An environment for modern phylogenetics and evolutionary analyses in R. Bioinformatics, 35(3), 526–528. 10.1093/bioinformatics/bty633

Parker, A. H., Zhang, P., Man, K. Y., Tirot, L., Rolfe, S. A., Smith, L. M., Wilkinson, S. W., & Ton, J. (2025). Contrasting epigenetic dynamics in chromosome arms and pericentromeres during ROS1-induced plant immune memory. 10.1101/2025.08.19.671022

Patro, R., Duggal, G., Love, M. I., Irizarry, R. A., & Kingsford, C. (2017). Salmon provides fast and bias-aware quantification of transcript expression. Nature Methods, 14(4), 417– 419. 10.1038/nmeth.4197

Pham, G. M., Hamilton, J. P., Wood, J. C., Burke, J. T., Zhao, H., Vaillancourt, B., Ou, S., Jiang, J., & Buell, C. R. (2020). Construction of a chromosome-scale long-read reference genome assembly for potato. GigaScience, 9(9), giaa100. 10.1093/gigascience/giaa100

R Core Team. (2026). R: a language and environment for statistical computing (Version 4.6.0) [Computer software]. R Foundation for Statistical Computing. 10.32614/R.manuals

Ramírez, F., Ryan, D. P., Grüning, B., Bhardwaj, V., Kilpert, F., Richter, A. S., Heyne, S., Dündar, F., & Manke, T. (2016). deepTools2: A next generation web server for deep-sequencing data analysis. Nucleic Acids Research, 44(W1), W160–W165. 10.1093/nar/gkw257

Reynoso, M. A., Kajala, K., Bajic, M., West, D. A., Pauluzzi, G., Yao, A. I., Hatch, K., Zumstein, K., Woodhouse, M., Rodriguez-Medina, J., Sinha, N., Brady, S. M., Deal, R. B., & Bailey-Serres, J. (2019). Evolutionary flexibility in flooding response circuitry in angiosperms. Science, 365(6459), 1291–1295. 10.1126/science.aax8862

Ritchie, M. E., Phipson, B., Wu, D., Hu, Y., Law, C. W., Shi, W., & Smyth, G. K. (2015). Limma powers differential expression analyses for RNA-sequencing and microarray studies. Nucleic Acids Research, 43(7), e47–e47. 10.1093/nar/gkv007

Rizhsky, L., Liang, H., & Mittler, R. (2002). The Combined Effect of Drought Stress and Heat Shock on Gene Expression in Tobacco. Plant Physiology, 130(3), 1143–1151. 10.1104/pp.006858

Rizhsky, L., Liang, H., Shuman, J., Shulaev, V., Davletova, S., & Mittler, R. (2004). When Defense Pathways Collide. The Response of Arabidopsis to a Combination of Drought and Heat Stress. Plant Physiology, 134(4), 1683–1696. 10.1104/pp.103.033431

Robinson, M. D., McCarthy, D. J., & Smyth, G. K. (2010). edgeR : a Bioconductor package for differential expression analysis of digital gene expression data. Bioinformatics, 26(1), 139–140. 10.1093/bioinformatics/btp616

Schep, A. N., Wu, B., Buenrostro, J. D., & Greenleaf, W. J. (2017). chromVAR: inferring transcription-factor-associated accessibility from single-cell epigenomic data. Nature Methods, 14(10), 975–978. 10.1038/nmeth.4401

Schwacke, R., Ponce-Soto, G. Y., Krause, K., Bolger, A. M., Arsova, B., Hallab, A., Gruden, K., Stitt, M., Bolger, M. E., & Usadel, B. (2019). MapMan4: A Refined Protein Classification and Annotation Framework Applicable to Multi-Omics Data Analysis. Molecular Plant, 12(6), 879–892. 10.1016/j.molp.2019.01.003

Shannon, P., Markiel, A., Ozier, O., Baliga, N. S., Wang, J. T., Ramage, D., Amin, N., Schwikowski, B., & Ideker, T. (2003). Cytoscape: A Software Environment for Integrated Models of Biomolecular Interaction Networks. Genome Research, 13(11), 2498–2504. 10.1101/gr.1239303

Soneson, C., Love, M. I., & Robinson, M. D. (2016). Differential analyses for RNA-seq: Transcript-level estimates improve gene-level inferences. F1000Research, 4, 1521. 10.12688/f1000research.7563.2

Subramanian, A., Tamayo, P., Mootha, V. K., Mukherjee, S., Ebert, B. L., Gillette, M. A., Paulovich, A., Pomeroy, S. L., Golub, T. R., Lander, E. S., & Mesirov, J. P. (2005). Gene set enrichment analysis: A knowledge-based approach for interpreting genome-wide expression profiles. Proceedings of the National Academy of Sciences, 102(43), 15545–15550. 10.1073/pnas.0506580102

Sullivan, A. M., Arsovski, A. A., Lempe, J., Bubb, K. L., Weirauch, M. T., Sabo, P. J., Sandstrom, R., Thurman, R. E., Neph, S., Reynolds, A. P., Stergachis, A. B., Vernot, B., Johnson, A. K., Haugen, E., Sullivan, S. T., Thompson, A., Neri, F. V., Weaver, M., Diegel, M., … Stamatoyannopoulos, J. A. (2014). Mapping and Dynamics of Regulatory DNA and Transcription Factor Networks in A. thaliana. Cell Reports, 8(6), 2015–2030. 10.1016/j.celrep.2014.08.019

Tamura, K., & Bono, H. (2022). Meta-Analysis of RNA Sequencing Data of Arabidopsis and Rice under Hypoxia. Life, 12(7), 1079. 10.3390/life12071079

Tournaire-Roux, C., Sutka, M., Javot, H., Gout, E., Gerbeau, P., Luu, D.-T., Bligny, R., & Maurel, C. (2003). Cytosolic pH regulates root water transport during anoxic stress through gating of aquaporins. Nature, 425(6956), 393–397. 10.1038/nature01853

Trapero-Mozos, A., Ducreux, L. J. M., Bita, C. E., Morris, W., Wiese, C., Morris, J. A., Paterson, C., Hedley, P. E., Hancock, R. D., & Taylor, M. (2018). A reversible light- and genotype-dependent acquired thermotolerance response protects the potato plant from damage due to excessive temperature. Planta, 247(6), 1377–1392. 10.1007/s00425-018-2874-1

Van Bel, M., Silvestri, F., Weitz, E. M., Kreft, L., Botzki, A., Coppens, F., & Vandepoele, K. (2022). PLAZA 5.0: Extending the scope and power of comparative and functional genomics in plants. Nucleic Acids Research, 50(D1), D1468–D1474. 10.1093/nar/gkab1024

van Berloo, R., Hutten, R. C. B., van Eck, H. J., & Visser, R. G. F. (2007). An Online Potato Pedigree Database Resource. Potato Research, 50(1), 45–57. 10.1007/s11540-007-9028-3

Vo, H. M., Charleston, M. A., Brodribb, T. J., & Sussmilch, F. C. (2025). Dehydration rapidly induces expression of NCED genes from a single subclade in diverse eudicots. Planta, 261(2), 46. 10.1007/s00425-025-04626-z

Wang, T.-Q., Peng, X.-R., Gao, H., Wei, L., Wang, S.-M., Wu, Y., & Li, Y. J. (2025). Chromatin accessibility and transcriptome analyses reveal stress-responsive regulatory elements in *Zygophyllum xanthoxylum*. Plant Physiology, 199(2), kiaf503. 10.1093/plphys/kiaf503

Wang, Z.-W., Liu, M., Sang, H.-T., Hou, S., Li, G., Tian, R.-M., Yang, Y.-Y., Xie, K., Wang, L., Li, N.-N., & Jia, K.-H. (2026). Salt priming coordinates transcriptional and epigenetic states for enhanced salt tolerance in mung bean (Vigna radiata). Communications Biology, 9(1), 1384. 10.1038/s42003-026-10574-5

Wei, X., Zhang, H., Ma, R., Zang, J., Wang, Y., & Yu, C. (2026). Heat stress responses of plants during adaptation to high temperature environments: A review of recent advances and insights. Plant Signaling & Behavior, 21(1), 2735211. 10.1080/15592324.2026.2735211

Wen, S., Jian, H., Shang, L., Kear, P. J., Zhang, M., Li, Y., Yuan, P., & Lyu, D. (2025). Comprehensive transcriptional regulatory networks in potato through chromatin accessibility and transcriptome under drought and salt stresses. The Plant Journal, 121(6), e70081. 10.1111/tpj.70081

Xu, S., Li, L., Luo, X., Chen, M., Tang, W., Zhan, L., Dai, Z., Lam, T. T., Guan, Y., & Yu, G. (2022). *Ggtree*: A serialized data object for visualization of a phylogenetic tree and annotation data. iMeta, 1(4), e56. 10.1002/imt2.56

Yeo, D. S. G., Eydam, J., Bruckmüller, J., Kauder, F., Lübeck, J., Kaier, A., Sonnewald, S., & Sonnewald, U. (2026). Genotype-dependent DNA methylation patterns are negatively associated with allelic variation rather than heat-induced gene expression in two contrasting potato genotypes. The Plant Journal, 125(2), e70690. 10.1111/tpj.70690

Yoshiba, Y., Kiyosue, T., Nakashima, K., Yamaguchi-Shinozaki, K., & Shinozaki, K. (1997). Regulation of Levels of Proline as an Osmolyte in Plants under Water Stress. Plant and Cell Physiology, 38(10), 1095–1102. 10.1093/oxfordjournals.pcp.a029093

Yu, G., Wang, L.-G., & He, Q.-Y. (2015). ChIPseeker: An R/Bioconductor package for ChIP peak annotation, comparison and visualization. Bioinformatics, 31(14), 2382–2383. 10.1093/bioinformatics/btv145

Zagorščak, M., Abdelhakim, L., Rodriguez-Granados, N. Y., Široká, J., Ghatak, A., Bleker, C., Blejec, A., Zrimec, J., Novák, O., Pěnčík, A., Baebler, Š., Perez Borroto, L., Schuy, C., Županič, A., Afjehi-Sadat, L., Wurzinger, B., Weckwerth, W., Pompe Novak, M., Knight, M. R., … Teige, M. (2025). Integration of multi-omics data and deep phenotyping provides insights into responses to single and combined abiotic stress in potato. Plant Physiology, 197(4), kiaf126. 10.1093/plphys/kiaf126

Zagorščak, M., Zrimec, J., Bleker, C., Nolte, N., Juteršek, M., Ramšak, Ž., Gruden, K., & Petek, M. (2024). Evidence-based unification of potato gene models with the UniTato collaborative genome browser. Frontiers in Plant Science, 15. 10.3389/fpls.2024.1352253

Zandalinas, S. I., & Mittler, R. (2022). Plant responses to multifactorial stress combination. New Phytologist, 234(4), 1161–1167. 10.1111/nph.18087

Zandalinas, S. I., Sengupta, S., Fritschi, F. B., Azad, R. K., Nechushtai, R., & Mittler, R. (2021). The impact of multifactorial stress combination on plant growth and survival. New Phytologist, 230(3), 1034–1048. 10.1111/nph.17232

Zhang, H., & Sonnewald, U. (2017). Differences and commonalities of plant responses to single and combined stresses. The Plant Journal, 90(5), 839–855. 10.1111/tpj.13557

Zhou, J., Wang, J., Zheng, Z., Fan, B., Yu, J.-Q., & Chen, Z. (2015). Characterization of the promoter and extended C-terminal domain of Arabidopsis WRKY33 and functional analysis of tomato WRKY33 homologues in plant stress responses. Journal of Experimental Botany, 66(15), 4567–4583. 10.1093/jxb/erv221

Zhou, Y., Zhang, Z., Bao, Z., Li, H., Lyu, Y., Zan, Y., Wu, Y., Cheng, L., Fang, Y., Wu, K., Zhang, J., Lyu, H., Lin, T., Gao, Q., Saha, S., Mueller, L., Fei, Z., Städler, T., Xu, S.,… Huang, S. (2022). Graph pangenome captures missing heritability and empowers tomato breeding. Nature, 606(7914), 527–534. 10.1038/s41586-022-04808-9

