## Supporting information for "Combined heat and drought stress alters chromatin accessibility and primes potato abscisic acid biosynthesis"

**This file includes:**
Supplementary methods and results
Suppl. Figs S1–S11
References

**Provided as a separate Excel workbook:**
Suppl. Table S1. Consensus peak set composition.
Suppl. Table S2. Numbers of differentially accessible regions (DARs).
Suppl. Table S3. Gene set enrichment analysis (GSEA) results.
Suppl. Table S4. Differential motif activity (chromVAR).
Suppl. Table S5. Differential accessibility and expression results (DARs and DEGs).
Suppl. Table S6. Priming experiment: sample descriptions.
Suppl. Table S7. Priming experiment: soil moisture.
Suppl. Table S8. Priming experiment: physiological traits.
Suppl. Table S9. Drought memory classes.
Suppl. Table S10. MapMan enrichment of drought memory classes.
Suppl. Table S11. Priming experiment: differentially expressed genes in cv. Désirée.
Suppl. Table S12. Priming experiment: regulatory network.
Suppl. Table S13. Transactivation assay.

### SUPPLEMENTARY METHODS AND RESULTS

#### Soil moisture

Soil moisture content in the priming experiment was measured as the average of five replicate measurements taken from the growing substrate using gravimetric analysis. Percentage soil moisture was in the range 71-76% in the well-watered samples but fell to as low as 52% in the samples subjected to stress (Suppl. Fig. S8).

#### Relative Water Content

Relative Water Content (RWC) was measured on fully expanded upper third or fourth leaves from the priming experiment. Three biological replicates were collected from separate plants. The fresh weight of harvested leaflets was immediately measured prior to incubation in distilled water overnight at room temperature. Excess water was removed by blotting with tissue paper and the turgid weight was recorded. Turgid leaf samples were first dried in a microwave oven at 500 W power for 10 min and then in a drying oven at 95 ℃ for 2-3 hr. Finally, dried leaf sample weights were recorded.

RWC values of genotypes were calculated using the following equation:

$$\mathrm{RWC}\ (\%) = \frac{\text{Fresh weight} - \text{Dry weight}}{\text{Turgid weight} - \text{Dry weight}} \times 100$$

#### Photosynthetic and stomatal parameters

Photosynthetic rate, stomatal conductance, and internal $CO_2$ level were measured on fully expanded upper third or fourth leaves with a CIRAS-II portable photosynthesis system using a built-in light source. Photosynthetically active radiation (PAR) was maintained at 1500 $\mu mol\ m^{-2}s^{-1}$ and $CO_2$ concentration was maintained at 400 $\mu mol\ mol^{-1}$. Measurements were performed on nine biological replicates.

#### Statistical analysis and visualisation

Differences between temperature treatments within each treatment group were assessed using the Wilcoxon rank-sum test. P-values were adjusted for multiple comparisons using the Holm method. Differences between temperature treatments were considered statistically significant at adjusted $p < 0.05$. Results were visualized using ggplot2 v4.0.3.

For raw data used to plot Suppl. Figs S8 and S9, see Suppl. Table S7 (soil moisture) and Suppl. Table S8 (physiological traits).

### SUPPLEMENTARY FIGURES

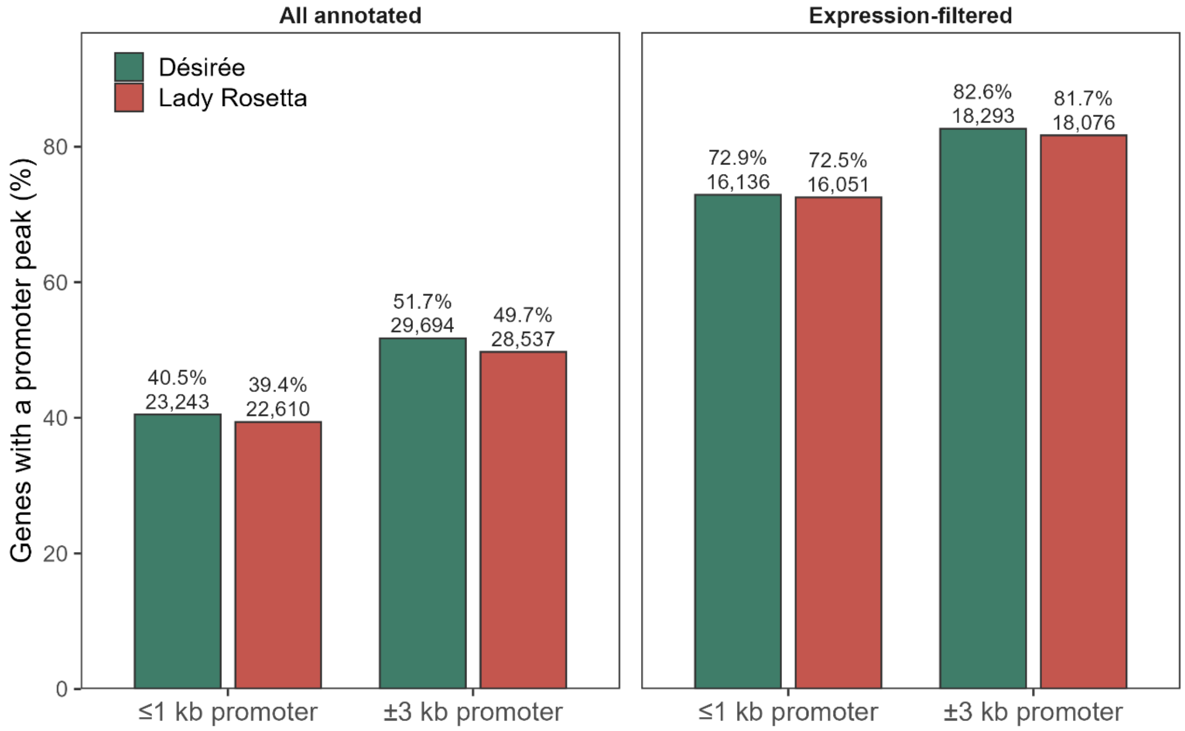


**Suppl. Fig. S1. Genes with an accessible promoter peak in unstressed plants.** Percentage of genes with at least one accessible chromatin region within 1 kb or 3 kb of the transcription start site in control plants of cvs. Désirée and Lady Rosetta. A region was counted if it was called in at least one of the three control harvest groups of the cultivar. Left, all 57,394 annotated genes; right, the 22,134 genes retained by the expression filter of the RNA-seq analysis. Bar labels give the percentage and the number of genes.

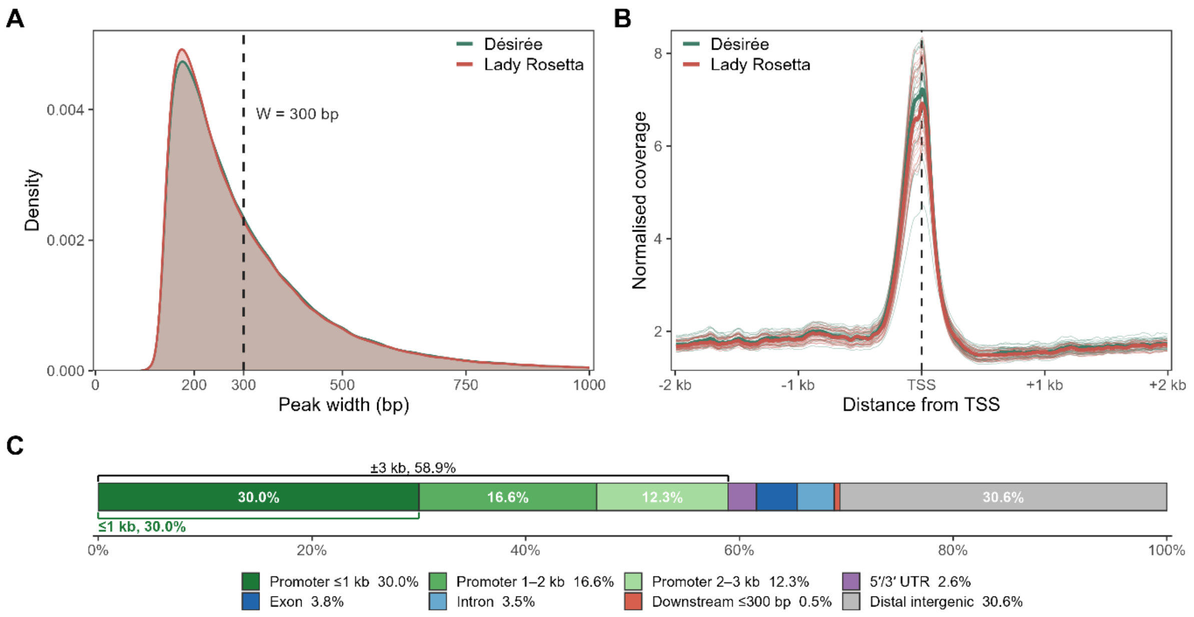


**Suppl. Fig. S2. Peak width distribution, accessibility around transcription start sites and genomic location of accessible chromatin regions. A)** Distribution of the widths of peaks called in each cultivar (Désirée, n = 612,342; Lady Rosetta, n = 608,617), pooled across treatment groups. Median peak widths were 259 bp and 256 bp, respectively. The dashed line marks the 300 bp window used to build the fixed-width consensus peak set. The x-axis is truncated at 1,000 bp; 1.6% (Désirée) and 1.5% (Lady Rosetta) of peaks are wider. **B)** Aggregated normalised read coverage around annotated transcription start sites (TSS), ±2 kb. Thin solid lines show individual libraries (n = 54; 27 Désirée and 27 Lady Rosetta) and thick solid lines the cultivar medians. Libraries are coloured by cultivar only. **C)** Genomic location of the consensus peak regions annotated against the UniTato gene models and shown as a proportional bar. Promoter regions are split by distance from the TSS. TSS, transcription start site; UTR, untranslated region; bp, base pairs; kb, kilobases.

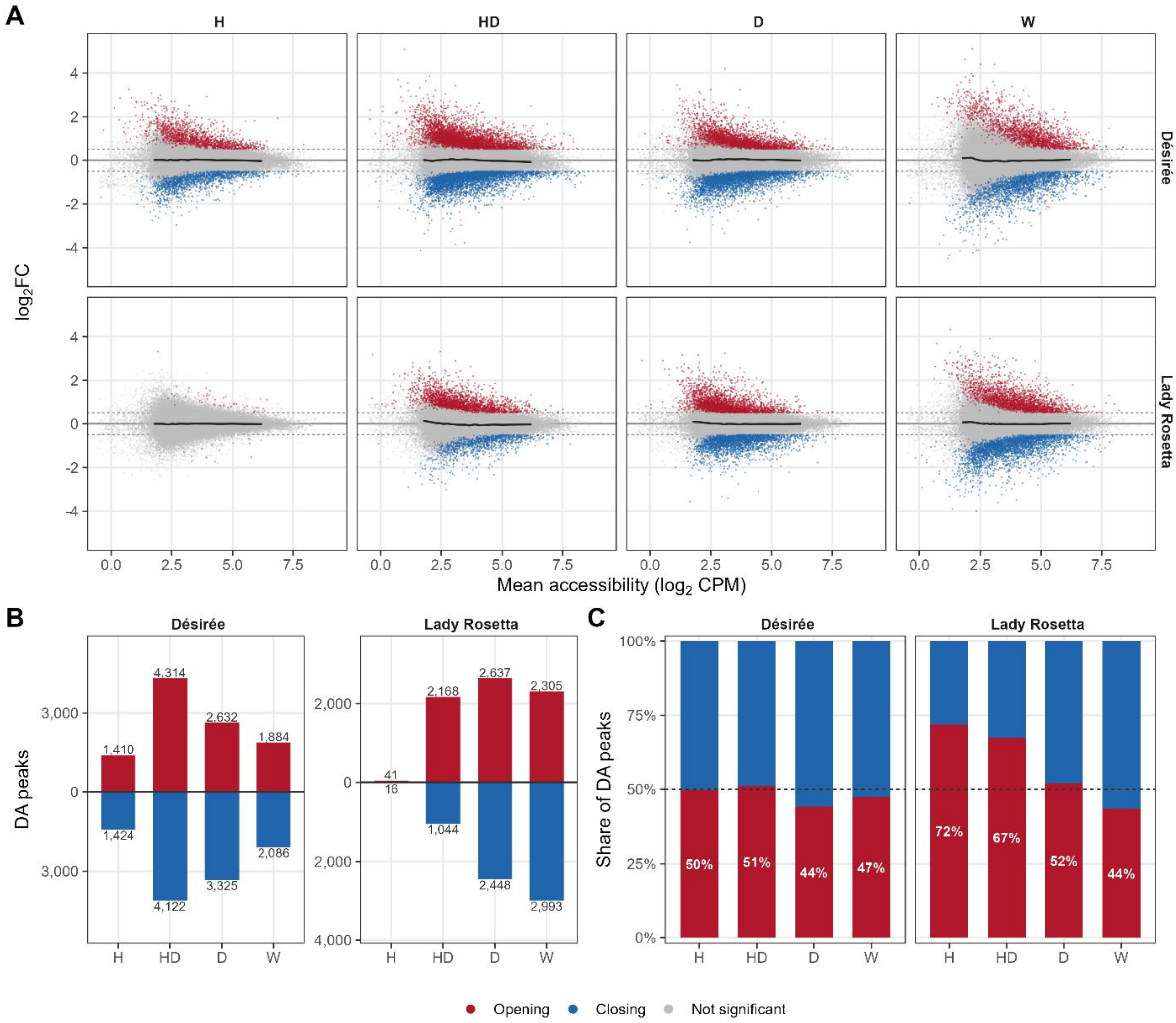


**Suppl. Fig. S3. Chromatin accessibility changes in both directions under every stress. A)** MA plots of the accessibility response in each stress comparison: $\log_2$ fold change ($\log_2$FC) against mean accessibility ($\log_2$ counts per million) for each of the 56,724 tested peaks. Differentially accessible peaks (FDR < 0.05 and $|\log_2FC| > 0.5$) are coloured by direction (red, opening; blue, closing) and all other peaks are grey. Dashed lines mark $|\log_2FC| = 0.5$. The dark line shows the median $\log_2$FC in 20 equal-count bins of mean accessibility. **B)** Number of differentially accessible peaks in each comparison, opening upwards and closing downwards, at the same thresholds. The vertical scales differ between cultivars. The number of differentially accessible peaks depends on the usable sequencing depth of the libraries compared, so bar heights are not comparable between stresses or cultivars. **C)** The same peaks as a percentage of all differentially accessible peaks in each comparison; the percentage of opening peaks is printed in the bar and the dashed line marks an even split. Heat in Lady Rosetta has only 57 differentially accessible peaks. FDR, false discovery rate; CPM, counts per million; H, heat; D, drought; W, waterlogging; HD, combined heat and drought.

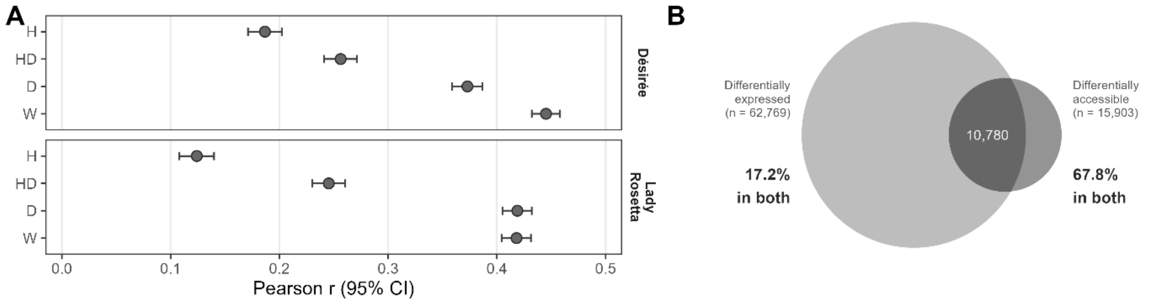


**Suppl. Fig. S4. Coupling and overlap between promoter accessibility and gene expression changes. A)** Correlation between the $\log_2$ fold changes of promoter accessibility and transcript abundance across all 14,716 genes with a tested promoter peak and an RNA-seq result, without selection on significance. Dots show Pearson's r coefficient with its 95% confidence interval for each stress and cultivar. **B)** Overlap between differentially expressed genes and genes with a differentially accessible promoter, pooled over the four stresses and both cultivars (gene-by-comparison tests). Circle areas are proportional to the number of tests in each set and the overlap area to their intersection. Accessibility is taken from the most significant promoter peak (within ±3 kb of the transcription start site) of each gene. Accessibility and expression comparisons were paired by cultivar, stress and sampling day, because the matched control differs between the two assays for drought and combined heat-drought. Significance was defined as FDR < 0.05 in each assay. The cultivars are shown side by side and are not compared statistically. FDR, false discovery rate; H, heat; D, drought; W, waterlogging; HD, combined heat and drought.

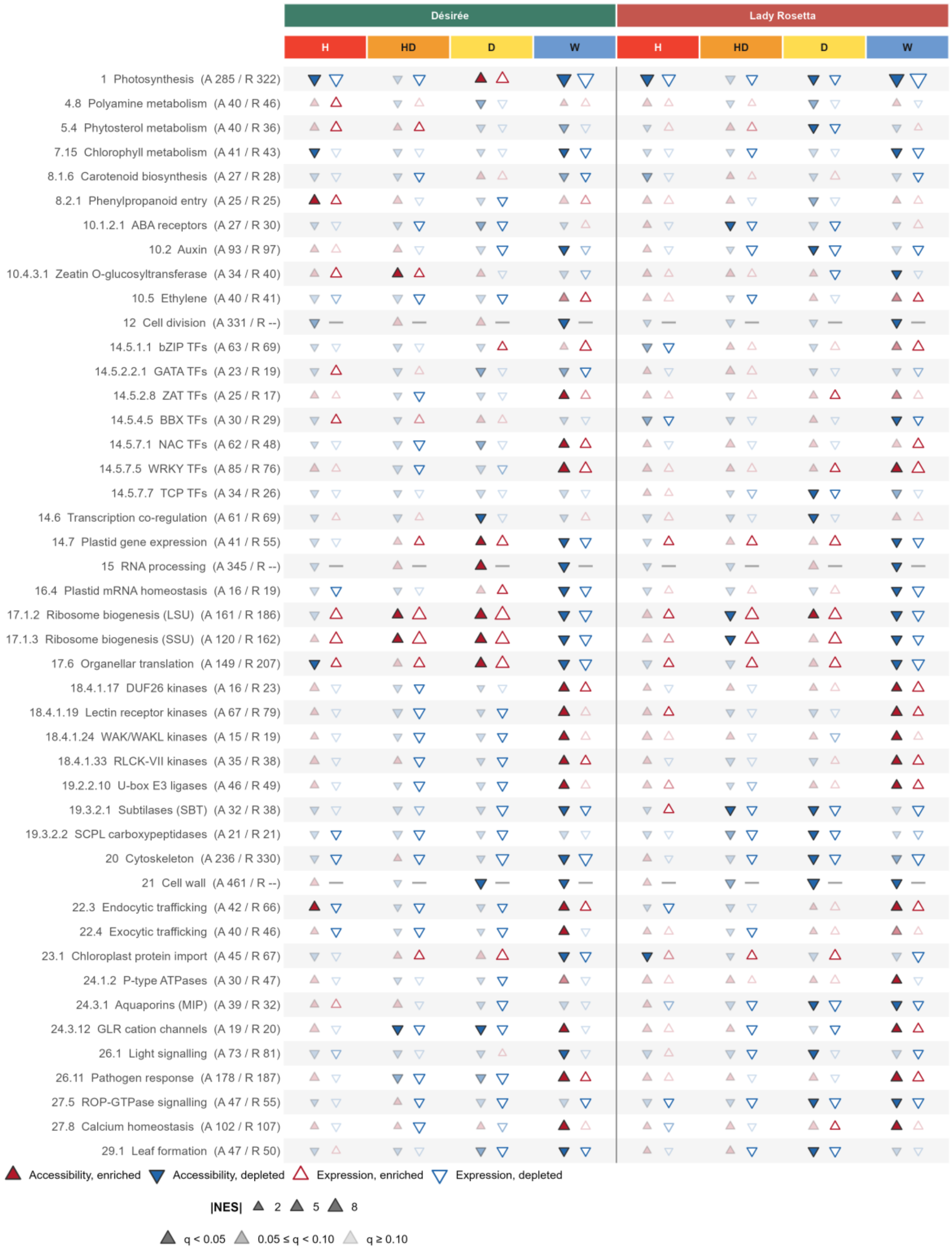


**Suppl. Fig. S5. MapMan gene set enrichment.** Enrichment of MapMan gene sets across all comparisons for every gene set with q < 0.10 in the accessibility dataset in at least 2 of the 8 comparisons, after nested gene sets were collapsed to a single representative gene set (45 gene sets); Fig. 2A shows only the selected 23. Each row label begins with the MapMan bin code, as in Suppl. Table S3, followed by the number of genes in the gene set for the accessibility (A) and expression (R) datasets. Filled and open triangles indicate enrichment in the accessibility and expression datasets, respectively; triangle orientation indicates increased or decreased accessibility or expression, and triangle size represents the absolute value of the normalised enrichment score (NES). Opacity indicates q value: <0.05, 0.05–0.10 and >0.10 correspond to 1, 0.5 and 0.22, respectively. A dash marks a gene set without an expression result, because it exceeds the maximum gene-set size in the expression dataset. Accessibility of the ribosome biogenesis gene sets (17.1.2 and 17.1.3) follows library quality (fraction of reads in peaks) rather than treatment and should therefore be interpreted with caution. Accessibility was tested only for genes with a promoter peak, so the accessibility and expression results rest on different gene sets and are compared by direction rather than magnitude. H, heat; D, drought; W, waterlogging; HD, combined heat and drought; NES, normalised enrichment score.

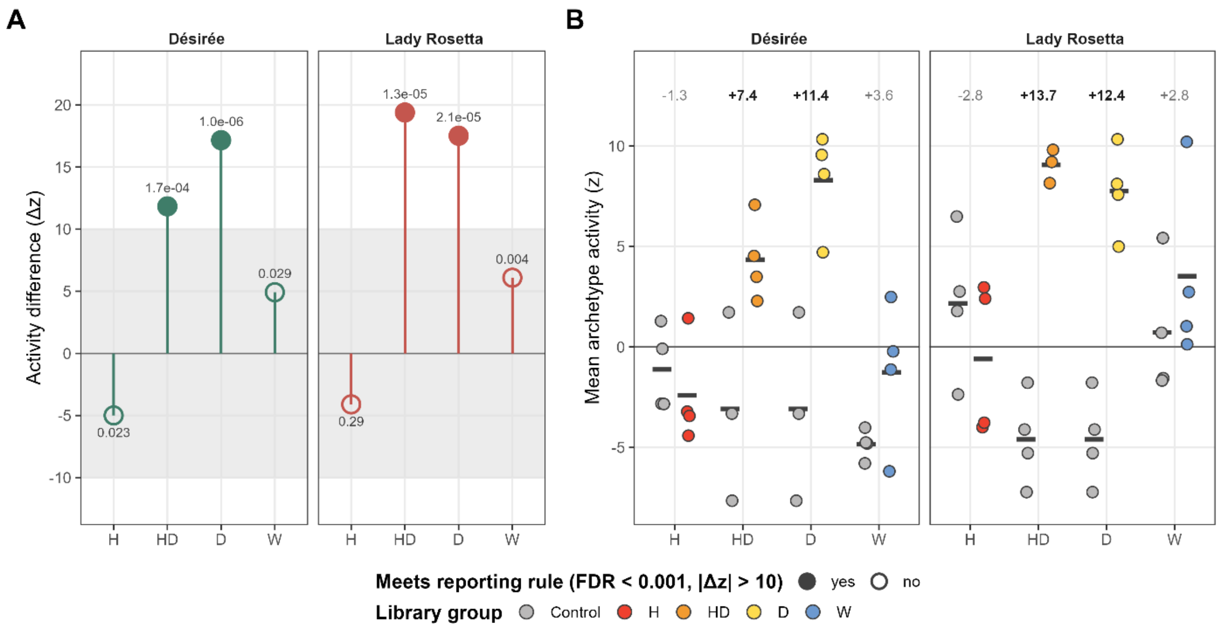


**Suppl. Fig. S6. Activity of the ABRE-type bZIP motif archetype across stresses. A)** Differential chromVAR activity of the ABRE-type bZIP motif archetype (JASPAR archetype JC_014, grouping 17 motif models at a correlation threshold of 0.70) in each stress compared with its matched control, given as the difference in activity (Δz; quantile-normalised chromVAR deviation z-scores). Filled points meet the reporting threshold (FDR < 0.001 and |Δz| > 10) and open points do not; the grey band spans |Δz| ≤ 10 and the FDR is printed next to each point. **B)** Archetype activity in individual libraries (mean z over its 17 motif models), with each stress group plotted next to its matched control group (C1 for waterlogging, C2 for heat and C3 for drought and heat-drought; the C3 group is shown twice per cultivar because drought and heat-drought share it). Horizontal bars indicate group means. The number above each pair is the stress minus control difference averaged over the 17 motif models, in bold where the comparison meets the reporting threshold. Motif activity was calculated with chromVAR on GC- and accessibility-matched background peaks, quantile-normalised and tested with limma separately in each cultivar. FDR values are Benjamini–Hochberg adjusted p-values across the 808 motif models of the JASPAR plants collection. C, control; H, heat; D, drought; W, waterlogging; HD, combined heat and drought; FDR, false discovery rate.

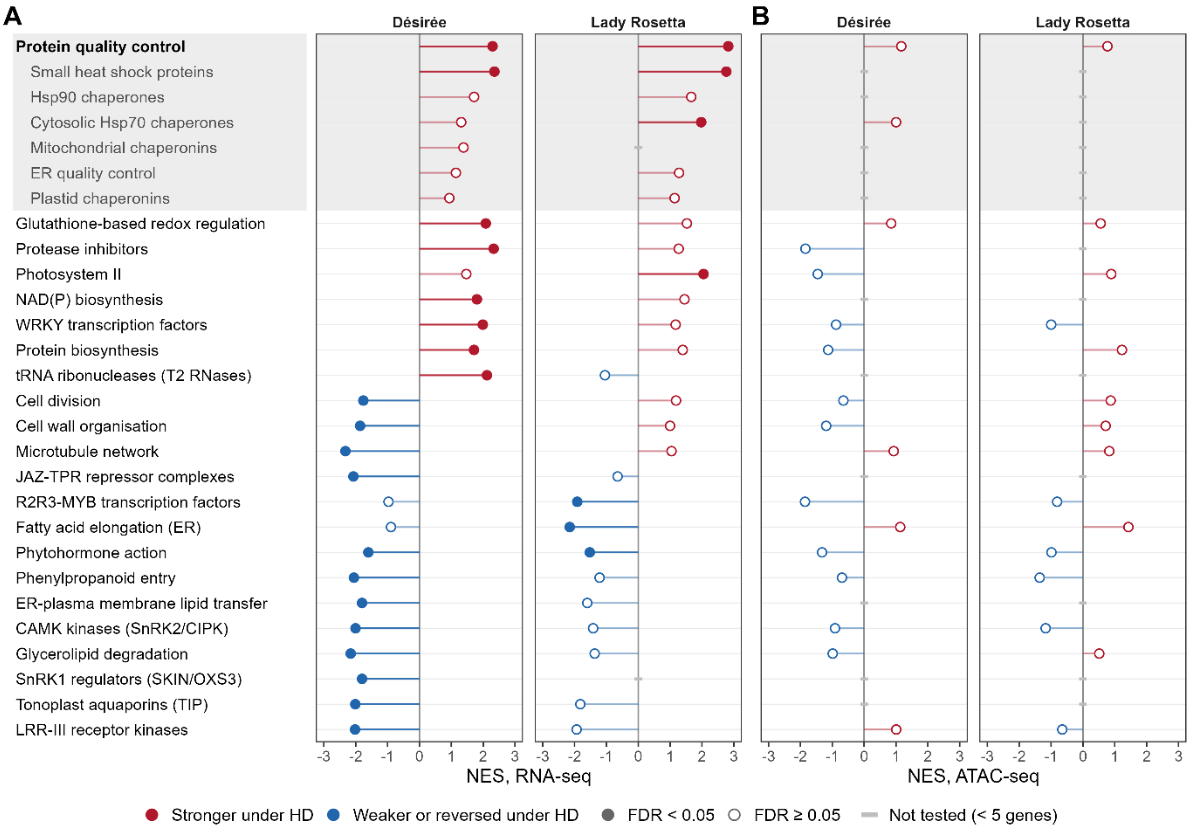


**Suppl. Fig. S7. Functional categories enriched among genes whose response differs between drought and combined heat and drought. A)** MapMan bins enriched along the shift in transcript response (RNA-seq) between drought and heat-drought in Désirée and Lady Rosetta. Points show the normalised enrichment score (NES) of each bin: red for a response stronger under heat-drought than under drought, blue for a weaker or reversed one; filled points are significant and open points are not. Rows are ordered by the mean NES of the two cultivars. Protein quality control subcategories are shaded. **B)** The same bins tested on promoter accessibility (ATAC-seq). A grey dash marks a bin not tested in that cultivar. D, drought; HD, combined heat and drought.

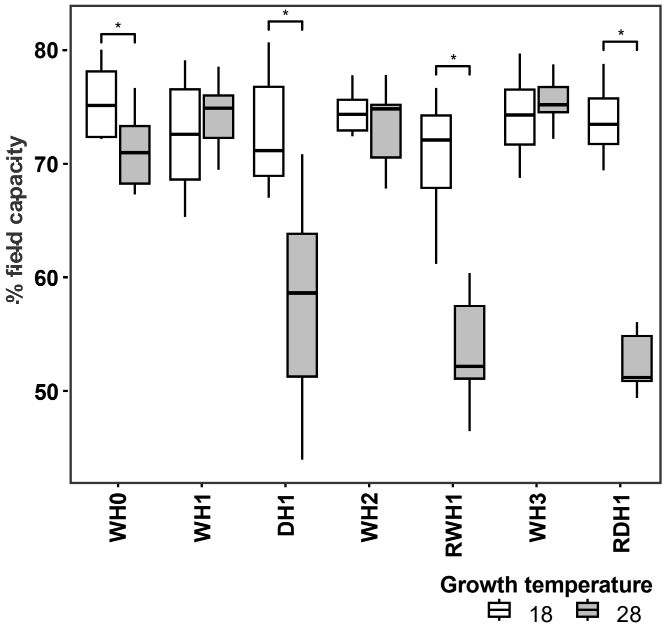


**Suppl. Fig. S8. Soil moisture in the priming experiment.** Box plots show soil moisture (%) of the growing substrate of plants grown at 18 °C and 28 °C at the sampling points of the experiment with two drought stress cycles. Boxes represent the interquartile range (IQR), horizontal lines indicate medians and whiskers extend to 1.5 × IQR. Asterisks denote significant differences between temperature treatments within each treatment group (Wilcoxon rank-sum test, Holm-adjusted $p < 0.05$). For sample details refer to Suppl. Table S6.

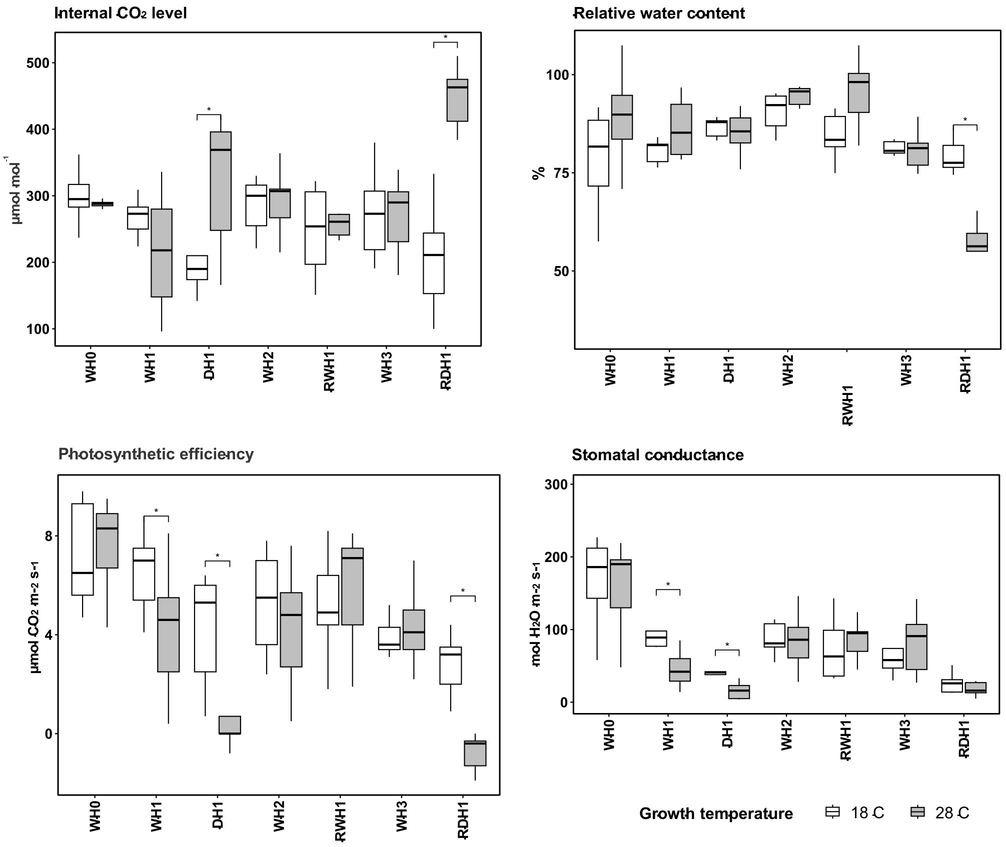


**Suppl. Fig. S9. Impact of stresses on leaf physiology.** Box plots show internal $CO_2$ level (µmol mol$^{-1}$), stomatal conductance, photosynthetic rate and relative water content (RWC, %) of leaves from plants grown at 18 °C and 28 °C at the sampling points of the experiment with two drought stress cycles. Boxes represent the interquartile range (IQR), horizontal lines indicate medians and whiskers extend to 1.5 × IQR. Asterisks denote significant differences between temperature treatments within each treatment group (Wilcoxon rank-sum test, Holm-adjusted $p < 0.05$). For sample details refer to Suppl. Table S6.

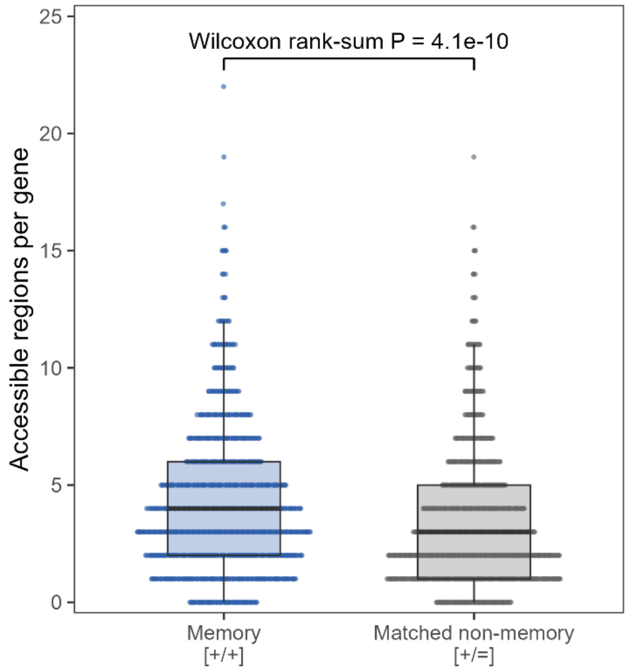


**Suppl. Fig. S10. Drought-memory genes carry more accessible regions.** Accessible regions per gene, for drought-memory genes (induced in the first drought and more strongly in the second; Ding et al. class [+/+]) and their matched non-memory genes (induced equally in both droughts; [+/=]; matching Fig. 4B). Boxes show the median and interquartile range (IQR), whiskers extend to 1.5 × IQR and points are individual genes. Median 4 (IQR 2–6) against 3 (IQR 1–5) regions per gene; Wilcoxon rank-sum test, $P = 4.1 \times 10^{-10}$.

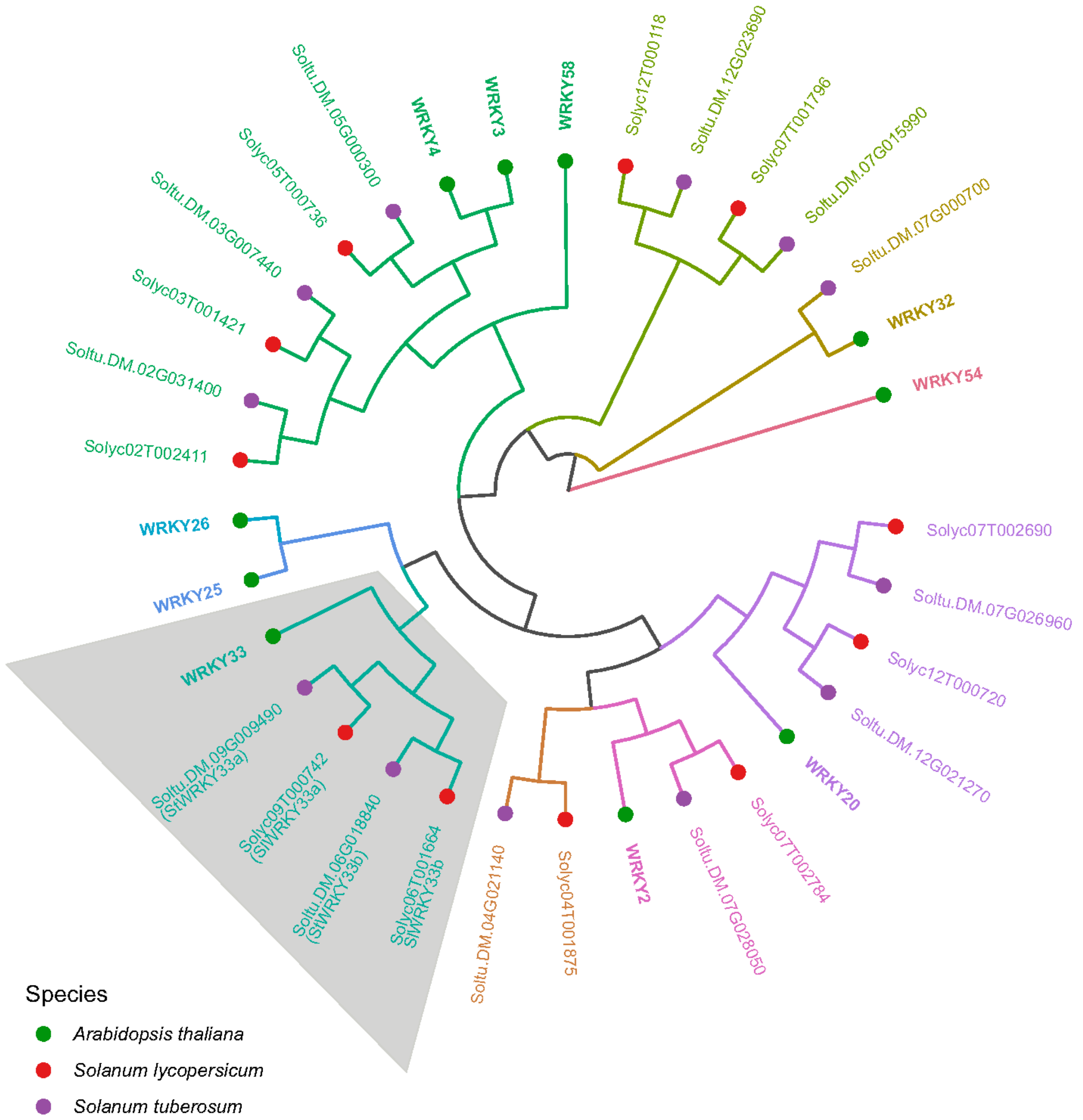


**Suppl. Fig. S11. Phylogeny of group I WRKY transcription factors.** Phylogenetic relationships of group I WRKY transcription factors from arabidopsis, potato and tomato. Branch and label colours indicate clusters, tip symbols indicate species, and Arabidopsis genes are shown in bold. The grey area marks the WRKY33 clade, which contains two WRKY33 orthologues in both Solanaceae species, located on chromosomes 9 and 6. The tomato genes Solyc09T000742 and Solyc06T001664 (genome annotation ITAG5.0) (Y. Zhou et al., 2022) correspond to Solyc09g014990 and Solyc06g066370, described as *SlWRKY33a* and *SlWRKY33b* by J. Zhou et al. (2015); the potato orthologues were named *StWRKY33a* (Soltu.DM.09G009490) and *StWRKY33b* (Soltu.DM.06G018840) accordingly. The tree is drawn as a cladogram therefore branch lengths are not to scale. The tree was built by neighbour joining from pairwise protein distances (1 − sequence identity) and rooted on Arabidopsis WRKY54 (group III).

### REFERENCES


Zhou, J., Wang, J., Zheng, Z., Fan, B., Yu, J.-Q., & Chen, Z. (2015). Characterization of the promoter and extended C-terminal domain of Arabidopsis WRKY33 and functional analysis of tomato WRKY33 homologues in plant stress responses. *Journal of Experimental Botany*, *66*(15), 4567–4583. https://doi.org/10.1093/jxb/erv221

Zhou, Y., Zhang, Z., Bao, Z., Li, H., Lyu, Y., Zan, Y., Wu, Y., Cheng, L., Fang, Y., Wu, K., Zhang, J., Lyu, H., Lin, T., Gao, Q., Saha, S., Mueller, L., Fei, Z., Städler, T., Xu, S., … Huang, S. (2022). Graph pangenome captures missing heritability and empowers tomato breeding. *Nature*, *606*(7914), 527–534. https://doi.org/10.1038/s41586-022-04808-9
